# Spatial Gene Set Enrichment Analysis with Applications to Spatially Resolved Transcriptomic Data

**DOI:** 10.64898/2026.06.01.729158

**Authors:** Zizhao Xie, Yanghong Guo, Qiwei Li, Ying Ma

**Author notes:** Correspondence to: Qiwei Li and Ying Ma (ying).

## Abstract

Spatially resolved transcriptomics enables the systematic characterization of spatial gene expression variation across tissue sections. Spatially variable genes within the same biological pathway often exhibit similar spatial expression patterns, reflecting shared biological functions and tissue organization. However, existing gene set enrichment analysis methods typically ignore this spatial dependence, which may reduce power to detect spatially organized pathways and limit the interpretability of pathway-level findings. To address this limitation, we propose spaGSE, a Bayesian hierarchical model for spatial pathway enrichment analysis that integrates genelevel summary statistics from spatial expression analysis with predefined gene set annotations. spaGSE models latent spatially variable gene signals through a Gaussian mixture framework and links spatial variation to gene set membership using logistic regression. To support robust and interpretable inference, we impose a spike-and-slab prior on the enrichment coefficient. Through simulation studies and analyses of four public SRT datasets, we show that spaGSE is scalable and achieves higher power while maintaining false positive rate control compared with existing approaches. In real-data applications, spaGSE identifies biologically relevant pathways with coordinated spatial organization across cancer and developmental tissues, demonstrating the value of incorporating spatial information into pathway-level inference for spatial transcriptomics.

## 1 Introduction

### 1.1 Motivation

Gene expression profiling has advanced from bulk RNA sequencing (RNA-seq) to single-cell RNA sequencing (scRNA-seq), and most recently, spatially resolved transcriptomics (SRT). Bulk RNA-seq provides a global measurement of transcriptomic activity across a tissue or cell population, but it averages the gene expression over mixed cell types and therefore cannot resolve cellular heterogeneity or spatial organization. scRNA-seq substantially improves resolution by profiling gene expression at the individual cell level, enabling the characterization of diverse cell populations and cell states. However, because scRNA-seq dissociates cells from their native tissue context, spatial information is lost, limiting its ability to characterize tissue architecture and spatially localized biological processes. SRT technologies address this limitation by measuring gene expression together with spatial coordinates, providing a direct view of how molecular activity is organized within tissues [Moffitt et al., 2022, Jain and Eadon, 2024]. These technologies have been widely applied to study tissue organization, cellular interactions, and spatially localized biological processes across diverse systems, including embryonic development, cancer, and neurodegenerative diseases [Moses and Pachter, 2022].

A central goal in transcriptomic data analysis is to move beyond individual genes and identify coordinated biological programs operating at the pathway or gene set level. Gene set enrichment analysis (GSEA) addresses this goal by determining whether predefined or curated sets of functionally related genes show coordinated evidence of biological activity. In conventional transcriptomic analyses, GSEA is often applied to ranked gene-level statistics, such as differential expression statistics, association statistics, or other gene-level summary measures. For SRT data, an analogous strategy is to first summarize spatial evidence at the gene level, for example using statistics from spatially variable gene (SVG) detection methods [Svensson et al., 2018], and then aggregate these gene-level signals to the pathway level. However, most existing GSEA methods were developed for bulk or scRNA-seq data and are not designed to perform pathway-level inference from spatially informed gene-level evidence. Hence, when applied to SRT data, these methods may have limited power and interpretability because they do not model how gene-level spatial evidence accumulates within a pathway, nor do they account for uncertainty in whether individual genes truly contribute to pathway-level spatial activity (see Section 1.2 for details).

This limitation is particularly important because pathway activity in SRT data may be reflected not only by the magnitude of gene-level spatial evidence, but also by coordinated spatial organization among functionally related genes. In SRT, genes within the same biological gene set or pathway may exhibit similar spatial expression patterns, reflecting shared functional roles, localized tissue structures, or region-specific biological processes. Empirically, genes identified with strong spatial evidence by methods such as SPARK [Sun et al., 2020] often display coherent spatial expression patterns when they belong to the same biological pathway, whereas genes outside the pathway tend to show distinct or less structured spatial patterns. This suggests that pathway-level signals in SRT data are characterized not only by enrichment of gene-level statistics, but also by coherence in their spatial organization. Therefore, simply applying conventional enrichment methods to gene-level spatial statistics may miss important pathway-level spatial structure.

We illustrate this phenomenon using a motivating example from a human pancreatic ductal adenocarcinoma (PDAC) dataset [Moncada et al., 2020], which is analyzed in detail in Section 5.1. Figure 1 summarizes the key motivation of developing spaGSE. For example, several SVGs identified by SPARK, including *ITGB4*, *MYL9*, and *TRIM29*, belong to the same pancreatic cancer-related gene set (GRUETZMANN PANCREATIC CANCER UP) and exhibited high expression localized to tumor regions (Figure 1a). These genes are known to be involved in processes such as cell adhesion, and epithelial tumor progression, which are central to PDAC development and invasion [Lv et al., 2022, Huang et al., 2024, Liu et al., 2026]. Their colocalized expression suggests coordinated pathway activity within the tumor microenvironment (TME). In contrast, SVGs outside this gene set, such as *CYP2S1*, *ABCC3*, and *AAMDC*, show more heterogeneous and less coordinated spatial patterns. Consistent with this observation, SVGs within this gene set show significantly stronger spatial autocorrelation than SVGs outside the set, with median Moran’s I values of 0.305 and 0.054 (*p <* 0.001 based on a Wilcoxon rank-sum test), respectively (see Supplementary Figure S1). Here, Moran’s I quantifies the degree of spatial autocorrelation, with larger values indicating greater similarity in expression among neighboring spatial locations [Chen, 2023]. Additional details and the corresponding distributional comparison boxplot are provided in Supplementary Section S2. This example also illustrates why existing GSEA tools may be insufficient for pathway analysis in SRT data (Figure 1b). When gene-level spatial statistics are analyzed using conventional GSEA tools, spatially coherent cancer-related pathways may fail to reach statistical significance, even when multiple member genes show tumor-localized spatial patterns. Conversely, enrichment results may include gene sets with limited spatial or disease-specific interpretability, because the analysis does not directly evaluate whether the pathway-level signal corresponds to coherent organization in tissue space. These observations suggest that pathway-level spatial analysis should account for both gene-level spatial evidence and the shared biological structure encoded by gene set membership.

**Figure 1:**
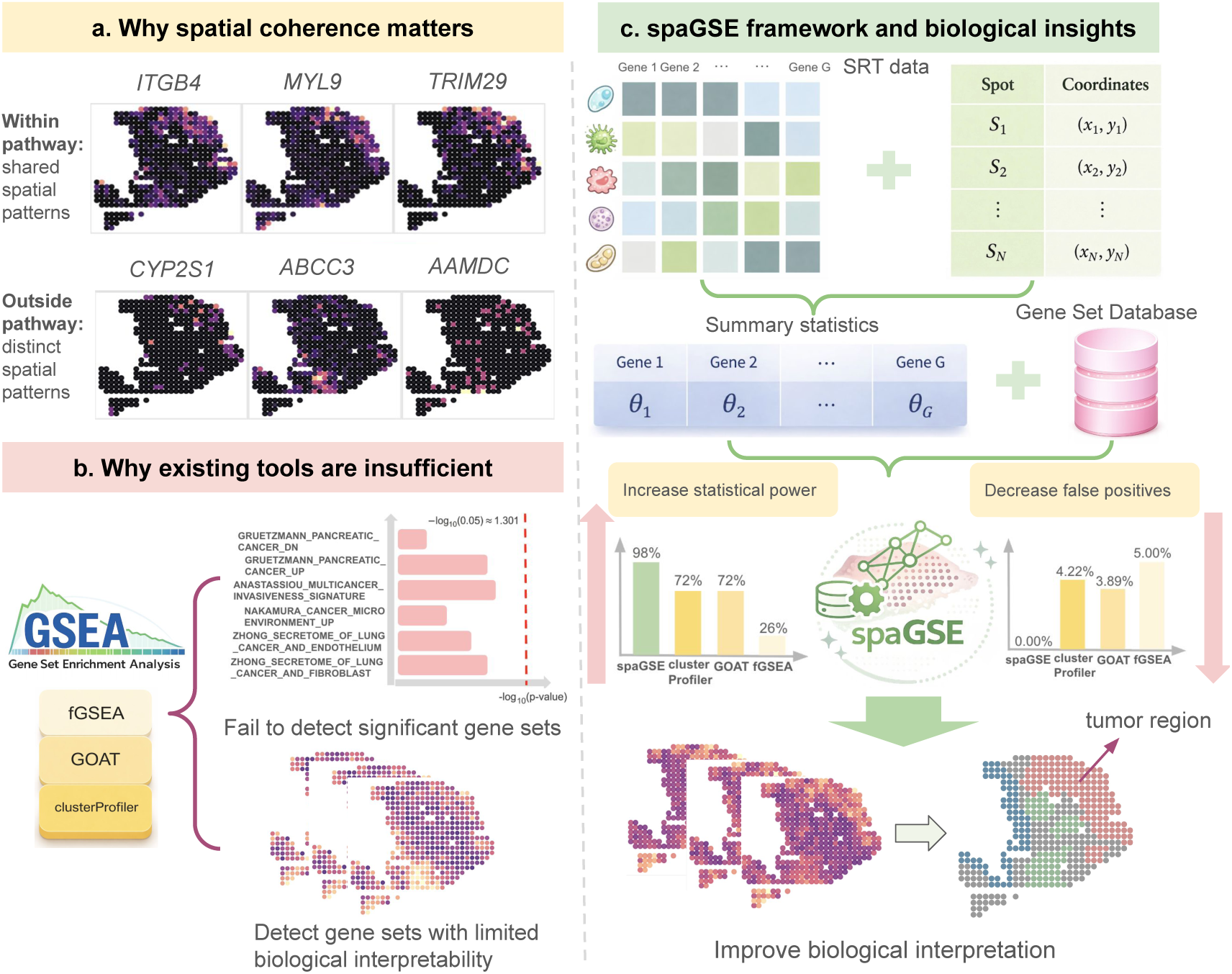
Schematic overview of the motivation and framework of spaGSE. Genes within the same pathway tend to exhibit shared spatial patterns, while genes outside the pathway show more distinct spatial variation. In the motivating example, SVGs within the target gene set show stronger spatial autocorrelation than SVGs outside the set, with mean Moran’s I values of 0.128 and 0.079, respectively; this difference is significant based on a Wilcoxon rank-sum test (*p <* 0.001). Existing GSE tools may therefore miss statistically significant and biologically meaningful gene sets when spatial information is ignored. spaGSE addresses this limitation by integrating spatial summary statistics and gene set information to improve statistical power, reduce false positives, and enhance biological interpretation.

Together, these observations indicate that pathway-level aggregation of spatial signals can reveal structured biological organization that is not fully captured by individual gene-level analyses or by conventional GSEA applied to spatial summary statistics. This motivates a joint probabilistic framework that integrates gene-level spatial evidence with pathway membership information, accounts for uncertainty in latent gene-level spatial relevance, and borrows information across genes to improve the detection and interpretation of biologically meaningful spatial programs from SRT data (Figure 1c).

### 1.2 Related Work

GSEA provides a pathway-level framework for interpreting high-throughput genomic data by aggregating signals across functionally related genes. Compared to analyses that focus only on individual genes, GSEA improves interpretability by identifying coordinated biological processes, such as metabolic pathways, transcriptional programs, and signaling networks [Subramanian et al., 2005]. However, most existing GSEA methods were developed for bulk and scRNA-seq data, including competitive approaches such as fGSEA [Korotkevich et al., 2016], clusterProfiler [Wu et al., 2021], and GOAT [Koopmans, 2024]. Specifically, fGSEA evaluates enrichment using a weighted Kolmogorov-Smirnov running sum statistic defined on a ranked gene list, with statistical significance assessed using a permutation test to construct a null distribution. clusterProfiler provides a flexible framework for enrichment analysis that includes both over-representation analysis and preranked GSEA. GOAT is a competitive approach for preranked gene lists that combines transformed gene rank scores with bootstrap null distributions to test whether genes in a given gene set tend to appear near the top of the ranking. These methods typically rely on ranked gene-level statistics and assess whether genes in a predefined set are enriched among the most significant signals. However, they treat genes as independent units and do not explicitly jointly model genes that share similar functions. In particular, these methods do not account for spatial gene expression patterns in SRT studies, which is a defining characteristic of SRT data, and may therefore reduce statistical power and fail to detect biologically meaningful pathways characterized by spatially coherent activity.

From a statistical perspective, borrowing information across related variables is known to substantially improve inference in high-dimensional settings. In many association studies, including genome-wide association studies (GWAS), it is well established that jointly modeling multiple variables within a Bayesian framework provides significant advantages over independent, univariate analyses [Zhou et al., 2013]. Even under simple composite likelihood formulations, where information is shared across predictors treated as conditionally independent, such approaches can substantially increase statistical power [Zhou et al., 2013]. In addition, Bayesian methods naturally provide uncertainty quantification for parameters of interest. These properties make Bayesian modeling particularly well suited for pathway-level analysis, where individual gene-level signals may be weak but become more informative when aggregated across functionally related genes.

To incorporate spatial information in pathway-level analysis, it is necessary to first quantify spatial variation at the gene level. In SRT studies, detecting SVGs is therefore a fundamental step for characterizing spatial expression heterogeneity, as it provides the primary evidence used in downstream pathway enrichment analysis [Li et al., 2022]. A wide range of computational methods have been developed for SVG detection, including Euclidean-distance-based and graph-based methods [Yan et al., 2025]. Euclidean-distance-based methods, such as SpatialDE [Svensson et al., 2018], Trendsceek [Edsgärd et al., 2018], SPARK [Sun et al., 2020], SPARK-X [Zhu et al., 2021], and BOOST-GP [Li et al., 2021], model spatial variation directly from spot coordinates. In contrast, graph-based frameworks such as Hotspot [DeTomaso and Yosef, 2021], BOOST-MI [Jiang et al., 2022], SpaGFT [Chang et al., 2024] rely on neighborhood graphs to capture local spatial structure, focusing on the connections between spots rather than their Euclidean distances. These methods differ in how they model spatial relationships, either through explicit coordinate-based kernels or neighborhood graph structures. Although these methods differ in their modeling strategies, they share a common output: gene-level summary statistics that quantify spatial variation. Benchmarking studies [Adhikari et al., 2024, Li et al., 2025] have shown that methods such as SPARK [Sun et al., 2020] and SPARK-X[Zhu et al., 2021] achieve strong performance in terms of type I error control and statistical power. In current practice, these summary statistics are subsequently used as inputs for pathway-level analysis. However, they are typically treated independently across genes, without accounting for shared structure within pathways or coordinated spatial patterns.

This limitation naturally motivates the use of joint modeling strategies that can borrow information across genes within a pathway. Such approaches have proved useful in related settings. For example, iDEA [Ma et al., 2020] is the only method that integrates differential expression analysis with GSEA into a unified framework to improve inference in scRNA-seq studies. However, extending these ideas to SRT is not straightforward. In particular, SRT data exhibit spatial dependence, and the gene-level inputs are typically summary statistics derived from SVG detection methods, such as test statistics or measures of spatial variation, rather than fold-change estimates with associated standard errors. As a result, existing integrative frameworks such as iDEA are not directly applicable in this setting. These considerations highlight the need for statistical methods that operate on gene-level spatial summary statistics while jointly modeling information across genes within pathways. Our proposed framework is designed to address this setting, enabling joint modeling of SVG-derived summary statistics to capture pathway-level spatial signals.

### 1.3 Contributions

We develop spaGSE, the first statistical framework specifically designed for spatial pathway enrichment analysis that integrates gene-level summary statistics from SVG detection with pathway membership information. By borrowing information across genes within a pathway, spaGSE improves the detection of enriched gene sets while providing uncertainty quantification for pathway enrichment effects through posterior inference.

Methodologically, spaGSE makes several contributions. First, it establishes a gene-level summary statistics-based modeling framework for pathway analysis in SRT, bridging the gap between SVG detection and pathway-level inference. Second, by modeling gene-level signals jointly within pathways, the approach accounts for dependence across genes, in contrast to existing enrichment methods that rely on marginal or independent summaries. Third, the Bayesian hierarchical formulation enables flexible modeling of pathway-level effects and uncertainty quantification, which is particularly important in settings where individual gene-level signals are weak but collectively informative. Finally, because the method operates directly on summary statistics rather than individual-level SRT data, it is computationally efficient, scalable, and broadly applicable across datasets and platforms, including large-scale studies with hundreds of thousands of spatial locations.

From an applied perspective, spaGSE provides a new framework for interpreting SRT data at the pathway level. Through comprehensive simulation studies and analyses of multiple SRT datasets across different platforms, we demonstrate that spaGSE improves statistical power and identifies biologically meaningful pathways that are closely linked to disease-relevant processes and tissue architecture, many of which are missed by existing approaches. These results highlight the importance of modeling pathway-level spatial coherence and illustrate how joint analysis of gene-level spatial signals can reveal biologically interpretable tissue architecture and disease microenvironments beyond gene-level analysis alone.

The remainder of this paper is organized as follows. Sections 2 and 3 introduce the proposed Bayesian hierarchical model and describe the Markov chain Monte Carlo (MCMC) algorithm used for posterior inference, respectively. Sections 4 and 5 evaluate the performance of spaGSE against existing approaches using both simulated datasets and four real SRT datasets across different platforms, respectively. Finally, Section 6 summarizes our findings and outlines potential directions for future research.

## 2 Model

We present spaGSE, a Bayesian hierarchical model for GSEA in SRT studies. spaGSE is designed to identify biological pathways whose member genes are more likely to exhibit spatial expression variation than genes outside the pathway. Rather than directly modeling the raw spatial gene expression count matrix for each gene set, spaGSE takes as input gene-level summary statistics (e.g., *z*-scores) generated from existing SVG detection methods, together with predefined gene set annotations. This summary-statistic formulation allows spaGSE to leverage a broad class of existing SVG detection methods while maintaining computational scalability for large SRT datasets. In addition, by performing GSEA on gene-level spatial evidence rather than directly on raw expression measurements, spaGSE reduces the complexity of pathway-level modeling and mitigates challenges arising from sparse counts, dropout events, platform-specific noise, and local spatial dependence.

An overview of the spaGSE workflow is illustrated in Figure 2. Starting from SRT count data and spatial coordinates, gene-level spatial evidence is first obtained using an SVG detection method (e.g., SPARK-X). These gene-level statistics are then incorporated into a Bayesian hierarchical model that introduces latent indicators for SVGs. Gene set membership is linked to the latent SVG gene status through a logistic regression model, allowing the enrichment coefficient to quantify whether genes in a given pathway have an increased probability of being SVGs. The model produces pathwaylevel statistical evidence, including Bayes factors (BFs) and credible intervals (CIs) for enrichment coefficients, which can be used to prioritize spatially enriched pathways for downstream biological interpretation. These prioritized pathways can then be mapped back to the tissue space to characterize how pathway-level spatial enrichment relates to local tissue architecture, disease-associated niches, and cellular microenvironmental organization.

**Figure 2:**
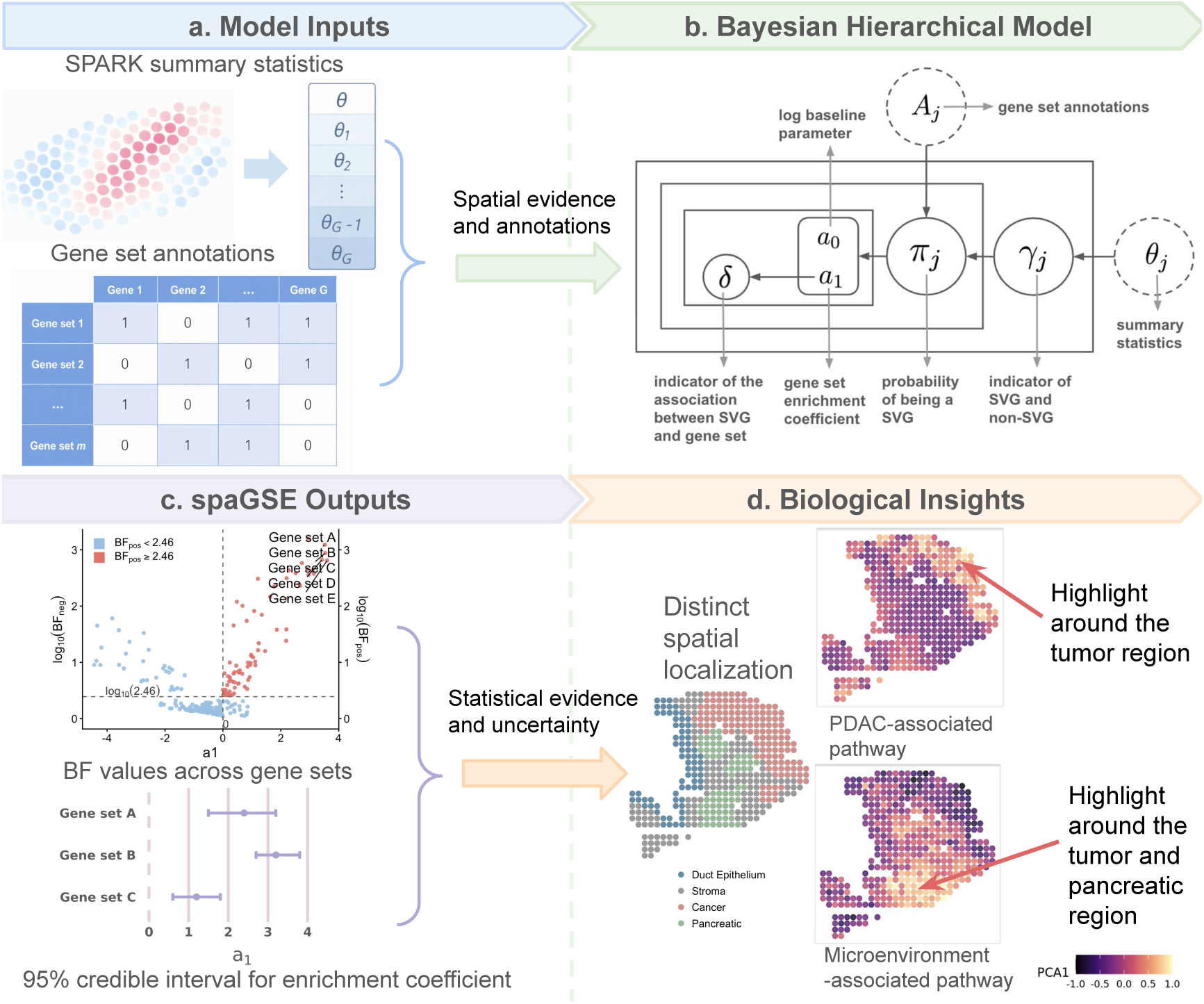
Overview of the spaGSE pipeline. Panel **a** shows the model inputs, including gene-level spatial evidence summarized from SPARK and gene set annotation information represented by a membership matrix. Panel **b** presents the Bayesian hierarchical model underlying spaGSE. Dashed circles denote observed inputs, whereas solid nodes denote latent variables and model parameters. The model links latent SVG status to gene set membership and quantifies pathway enrichment effects and uncertainty. Panel **c** illustrates the main statistical outputs of spaGSE, including Bayes factor (BF) values across gene sets and posterior credible intervals (CIs) for enrichment coefficients. Panel **d** highlights the biological insights provided by spaGSE, where enriched pathways show distinct spatial localization patterns. Example PDAC-related and microenvironment-associated pathways display coherent spatial activity around tumor and pancreatic regions.

### 2.1 Data Notation

Consider an SRT gene expression dataset measured at *N* spatial locations and across *J* genes. Let ***Y*** = [*y_ij_*]*_N×J_* denote the observed gene expression count matrix, where *y_ij_* ∈ N is the read count for gene *j* (*j* = 1*,…J*) observed at spatial location *i* (*i* = 1*,…N*). The corresponding spatial coordinates are denoted by ***T*** = [*t_id_*]*_N×_*_2_. Here, each row ***t****_i_* = (*t_i_*_1_*, t_i_*_2_) records the two-dimensional x and y-coordinates of spatial location *i*, allowing the model to capture the spatial relationships between locations in the tissue space. Although spatial coordinates are part of the original SRT data, they are not directly included in the spaGSE model. Instead, ***Y*** and ***T*** are used by upstream SVG detection methods to obtain gene-level spatial summary statistics, which are subsequently used as inputs to spaGSE for GSEA.

Before computing gene-level spatial summary statistics, each dataset is processed using standard quality-control procedures. Low-quality spatial locations and genes with very low expression are removed. Specifically, we applied the Seurat package (v4.1.1) [Hao et al., 2021] to filter genes detected in fewer than three spatial locations and locations with fewer than 100 detected genes. The filtered expression matrix and spatial coordinate matrix are then used as input to SVG detection methods.

Let *G_l_* denote the *l*-th predefined gene set, for *l* = 1*,…, L*. For a fixed gene set *G_l_*, we define the binary gene set membership indicator *A_jl_*, where *A_jl_* = 1 if gene *j* belongs to *G_l_* and *A_jl_* = 0 otherwise. The goal of spaGSE is to assess whether membership in *G_l_* is associated with an increased probability that a gene is an SVG exhibiting spatial variation. The key notation used throughout the proposed spaGSE model is summarized in Supplementary Table S1. This table provides definitions for the observed data, latent SVG indicators, enrichment parameters, prior specifications, MCMC quantities, and posterior inference metrics used in the model formulation and fitting.

### 2.2 Gene-level Spatial Summary Statistics

spaGSE uses gene-level spatial evidence obtained from existing SVG detection methods as input. As mentioned in Section 1.2, we primarily use SPARK and SPARK-X, two widely applied statistical methods for identifying SVGs. These methods model spatial dependence through kernel functions defined on the spatial coordinates and provide gene-level tests for spatial expression variation. For each gene *j*, let *p_j_* denote the *p*-value from the gene-level test of spatial variation from SPARK. We collect these values into the vector ***p*** = (*p*_1_*,…, p_J_*)*^T^* and transform them into nonnegative standard normal *z*-scores,

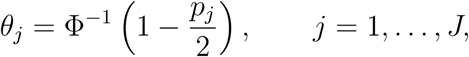

where Φ*^−^*^1^(·) denotes the inverse cumulative distribution function of the standard normal distribution. Larger values of *θ_j_* correspond to stronger evidence that gene *j* exhibits spatial variation. Let ***θ*** = (*θ*_1_*,…, θ_J_*)*^T^* denote the vector of transformed gene-level spatial statistics. The Bayesian enrichment model described below is fitted using ***θ*** together with the predefined gene set annotations.

### 2.3 Bayesian Hierarchical Model for GSEA

We introduce a Bayesian hierarchical model to characterize latent gene-level spatial signals and their associations with predefined gene sets. Since the transformed spatial summary statistics *θ_j_*’s represent the magnitude of gene-level spatial variation, we impose the non-negative support constraint *θ_j_* ∈ R^+^. To distinguish SVGs with spatial variation from the non-SVGs, we introduce a latent binary variable *γ_j_* ∈ {0, 1}, where *γ_j_* = 1 indicates that gene *j* is an SVG that exhibits a spatial expression pattern and *γ_j_*= 0 otherwise.

Conditional on *γ_j_*, we model *θ_j_* using a two-component Gaussian mixture distribution restricted to the nonnegative real line. Specifically,

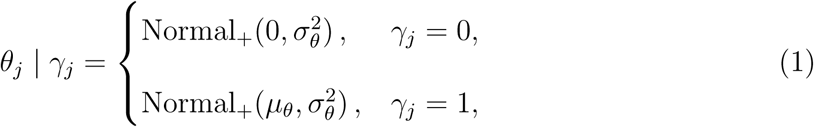

where Normal_+_(*µ_θ_, σ*^2^) denotes a normal distribution with mean parameter *µ_θ_* and variance parameter *σ*^2^ truncated to [0, ∞). This constraint is consistent with the interpretation of *θ_j_* as a nonnegative measure of spatial signal strength: non-SVGs are concentrated near zero, whereas SVGs are modeled by a component shifted toward larger positive values with *µ_θ_ >* 0. The latent indicator *γ_j_*is assigned a Bernoulli prior with probability *π_j_*,

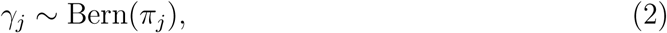

where *π_j_*denotes the prior probability that gene *j* shows spatial variation.

In our implementation, we set *µ_θ_*= 3 and *σ_θ_*= 1. Specifically, the choice *µ_θ_* = 3 reflects the average *z*-score observed among SVGs identified in real SRT datasets and represents strong gene-level evidence for spatial variation on the standard normal scale. We set *σ_θ_* = 1 to allow moderate variability around this alternative mean, consistent with the expected variability of *z*-scores for genes detected with approximately 90% statistical power by SPARK. In our simulation study, we evaluated the performance of spaGSE under different levels of gene-level variability by considering *σ_θ_* ∈ {0.2, 0.5}. We further conducted a sensitivity analysis for *µ_θ_* ∈ {1, 2, 3, 4}, with the results reported in the Supplementary Section S5. Overall, the results were consistent across different choices of *σ_θ_* and *µ_θ_*, indicating that the performance of spaGSE is robust to the specification of these prior parameters.

To connect gene-level spatial variation statistics with gene set membership, we model the prior probability *π_j_* through a logistic link:

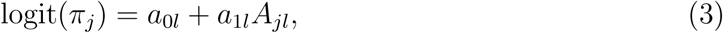

where *A_jl_*is a binary indicator of whether a gene *j* belongs to the gene set *l* of interest, the intercept term *a*_0_*_l_* represents the baseline log-odds that a gene outside the gene set is spatially variable, and the coefficient *a*_1_*_l_* is the gene set enrichment parameter. Because spaGSE analyzes each gene set separately, inference for multiple gene sets can be performed in parallel, which improves computational efficiency for large gene set collections. A positive value of *a*_1_*_l_* indicates that genes in the gene set have higher odds of being spatially variable than genes outside the gene set. Equivalently, *π_j_* is obtained by applying the inverse-logit transformation to *a*_0_*_l_* + *a*_1_*_l_A_jl_*:

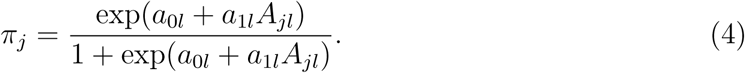

Under this formulation, pathway enrichment is assessed through inference on *a*_1_*_l_*. Evidence for positive enrichment corresponds to posterior support for *a*_1_*_l_ >* 0, whereas *a*_1_*_l_* = 0 corresponds to no difference in the probability of spatial variation between genes inside and outside the gene set. In practice, not all SVGs are associated with the gene set, therefore, to encourage sparse selection across gene sets, we specify a spike-and-slab prior for *a*_1_*_l_*:

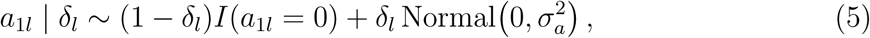

where *I*(·) is the indicator function, and hence, *I*(*a*_1*l*_ = 0) denotes a point mass at zero, and the binary indicator *δ_l_* determines whether the gene set has a nonzero enrichment effect.

When *δ_l_*= 0, the model sets *a*_1_*_l_* = 0, corresponding to no association between gene set membership and spatial variation. When *δ_l_* = 1, *a*_1_*_l_* follows a normal distribution, allowing the data to support either enrichment or depletion.

We consider a Bernoulli prior on *δ_l_*,

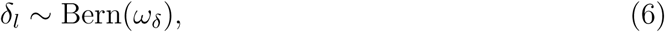

where *ω_δ_* denotes the prior probability that a gene set has a nonzero enrichment effect. To allow more flexibility, we further assign a conjugate beta hyperprior on *ω_δ_*,

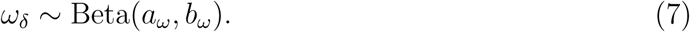

The hyperparameters *a_ω_*and *b_ω_*control the prior sparsity level across gene sets through the prior mean *a_ω_/*(*a_ω_* + *b_ω_*). This quantity represents the prior expected proportion of gene sets with nonzero enrichment effects. We set *a_ω_* + *b_ω_* = 2 so that the beta prior remains weakly informative while still encoding a preference for sparse pathway selection. In most applications, choosing a value of *a_ω_/*(*a_ω_* +*b_ω_*) ∈ [0.1, 0.2] encodes the prior expectation that approximately 10% to 20% of gene sets have nonzero enrichment effects. Unless otherwise stated, we set *a_ω_* = 0.4 and *b_ω_* = 1.6, which gives a prior mean of 0.2 and favors sparsity while allowing a small subset of gene sets to be associated with SVGs.

Finally, for *a*_0_*_l_* in Equation (3), we assign a weakly informative normal hyperprior to avoid imposing strong prior constraints on the magnitude or direction of the enrichment effects. Specifically, we set *a*_0_*_l_* ∼ N(0*, σ*^2^) and use relatively large values for *σ_a_* and *σ*_0_, such as 10. Here, *σ*^2^ controls the prior variability of the baseline log-odds, while *σ*^2^ controls the prior variability of the nonzero enrichment effect under the slab component.

## 3 Model Fitting

The model parameters consist of the baseline log-odds parameter *a*_0_*_l_*, the gene set enrichment coefficient *a*_1_*_l_*, the indicator *δ_l_* determining whether the gene set has a nonzero enrichment effect, and the indicator *γ_j_*indicating whether each gene is an SVG. Conditional on the latent spatial indicators, the full data likelihood for the gene-level summary statistics ***θ*** = (*θ*_1_*,…,θ_J_*)*^T^* is:

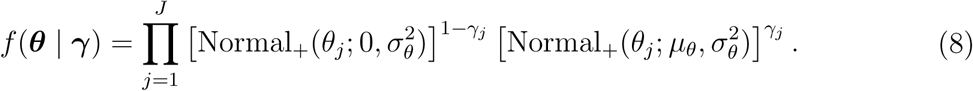

Combining the full data likelihood with the prior distributions specified in Section 2.3, the full posterior distribution is

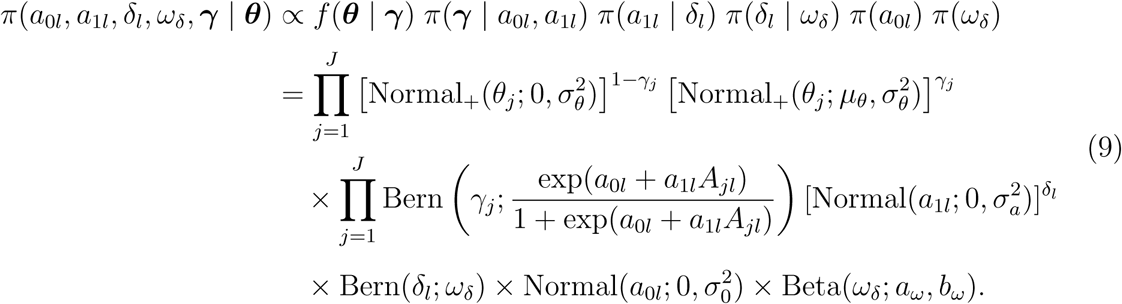

### 3.1 MCMC Algorithms

We implement the MCMC algorithm for posterior inference by updating each parameter with Metropolis-Hastings (MH) sampling. We update the parameters in each iteration following the steps below:

### Update

*γ_j_*: For each gene *j* = 1*,…,J*, we update the latent indicator *γ_j_* using a single-site flip proposal. Specifically, given the current value *γ_j_*, we propose *γ\** = 1 − *γ_j_*.

### The proposed value is accepted with probability min{1*, r*_MH_}, where

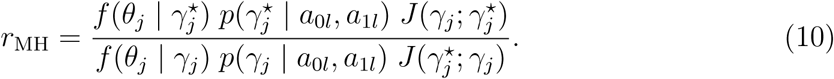

Here, *f* (*θ_j_* | *γ_j_*) and *f* (*θ_j_* | *γ\**) denote the likelihood contributions of the observed gene-level spatial summary statistic *θ_j_* under the current and proposed SVG indicators, respectively. The terms *p*(*γ_j_* | *a*_0_*_l_, a*_1_*_l_*) and *p*(*γ\** | *a*_0_*_l_, a*_1_*_l_*) denote the Bernoulli probabilities of the current and proposed indicators induced by logit(*π_j_*) = *a*_0_*_l_* + *a*_1_*_l_A_jl_*. Finally, *J* (*γ_j_*; *γ\**) and *J* (*γ\**; *γ_j_*) denote the forward and reverse proposal probabilities, respectively.

### Jointly update *δ_l_*and *a*_1_*_l_*

We jointly update the inclusion indicator *δ_l_* and the enrichment coefficient *a*_1_*_l_* using an *add-delete* MH algorithm. This step allows the sampler to move between the null model, in which *a*_1_*_l_* = 0, and the alternative model, in which *a*_1_*_l_* is estimated from the data.

For an *add* move, i.e. *δ_l_* = 0 → *δ_l_\** = 1, we propose *a\** from Normal(0,τ*_a_*^2^). For the *delete* move, i.e. *δ_l_* = 1 → *δ_l_\** = 0, we propose *a\** = 0. The proposed pair (*δ _l_^*^, a\**_1_*_l_*) is accepted with probability min{1*, r*_MH_}, where

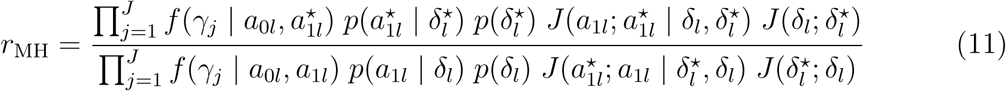

Here, *f* (*γ_j_* | *a*_0_*_l_, a*_1_*_l_*) and *f* (*γ_j_*| *a*_0_*_l_, a\**) denote the likelihood contributions of the latent SVG indicators under the current and proposed enrichment coefficients, respectively. Specifically, each *f* (*γ_j_*| *a*_0_*_l_, a*_1_*_l_*) is the Bernoulli probability induced by the logistic regression model for *π_j_*. The terms *p*(*a*_1_*_l_* | *δ_l_*) and *p*(*a\** | *δ\**) denote the spikeand-slab prior densities for the current and proposed enrichment coefficients, respectively, while *p*(*δ_l_*) and

*p*(*δ*_1_*^*^*) denote the prior probabilities of the current and proposed inclusion indicators. Finally, *J* (*a*_1_*_l_*; *a\** | *δ_l_, δ_l_ ^*^*), *J* (*a*_1_*_l_^*^*; *a*_1_*_l_* | *δ_l_*, δ_l_*), *J* (*δ_l_*; *δ_l_\**), and *J* (*δ_l_\**; *δ_l_*) denote the corresponding forward and reverse proposal probabilities for the add-delete move.

Update *a*_1_*_l_* when *δ_l_* = 1: When *δ_l_* = 1, we further update *a*_1_*_l_* within the alternative model using a random-walk MH (RWMH) step. We propose *a\** ∼ Normal(*a*_1_*_l_, τ* ^2^*/*2), and accept the proposed value with probability min{1*, r*_MH_}, where

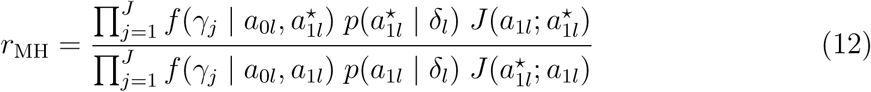

Here, *f* (*γ_j_* | *a*_0_*_l_, a*_1_*_l_*) denotes the Bernoulli probability mass function induced by the logistic regression model for *π_j_*, and *p*(*a*_1_*_l_* | *δ_l_*) denotes the slab prior density of *a*_1_*_l_* when

*δ_l_* = 1. The terms *J* (*a*_1_*_l_*; *a*_1_*_l_^*^*) and *J* (*a*_1_*_l_^*^*; *a*_1_*_l_*) denote the proposal densities for the reverse and forward moves, respectively.

Update *a*_0_*_l_*: We update the baseline log-odds parameter *a*_0_*_l_* using an RWMH algorithm. We propose a new *a\** from Normal(*a*_0_*_l_, τ* ^2^) and accept the proposed value with probability min{1*, r*_MH_}, where

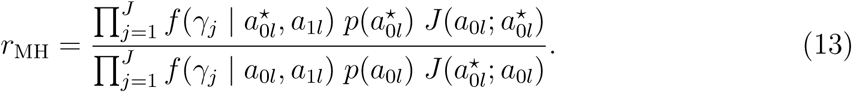

Here, *f* (*γ_j_* | *a*_0_*_l_, a*_1_*_l_*) and *f* (*γ_j_* | *a*_0l_, a*_1_*_l_*) denote the likelihood contributions of the latent SVG indicators under the current and proposed baseline log-odds parameters, respectively. Each *f* (*γ_j_* | *a*_0_*_l_, a*_1_*_l_*) is the Bernoulli probability induced by the logistic regression model for *π_j_*. The terms *p*(*a*_0_*_l_*) and *p*(*a\**) denote the prior densities of the current and proposed values of *a*_0_*_l_*. Finally, *J* (*a*_o_*_l_^*^*; *a*_0_*_l_*) and *J* (*a*_0_*_l_*; *a\**_o_*_l_*) denote the forward and reverse proposal probabilities, respectively.

Update

*ω_δ_*: Since *ω_δ_* has a conjugate Beta full conditional, the Gibbs sampler is

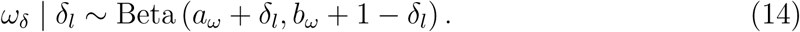

Because each gene set is analyzed separately, *ω_δ_* is updated conditional on the single inclusion indicator *δ_l_* for the current gene set. This update mainly reflects the prior sparsity assumption and provides the prior odds used in the BF calculation in Equation (15).

### 3.2 Posterior Inference

For posterior inference, our primary focus is the gene set enrichment coefficient *a*_1_*_l_*, which quantifies the association between gene set membership and the probability that a gene is spatially variable. After burn-in iterations, we use the retained MCMC samples to summarize the posterior distribution of *a*_1_*_l_* and evaluate evidence for pathway enrichment. Specifically, a positive value of *a*_1_*_l_* indicates that genes in the analyzed gene set have higher odds of being an SVG than genes outside the set. We therefore consider the null and alternative hypotheses as M_0_: *a*_1_*_l_* ≤ 0 and M_1_: *a*_1_*_l_ >* 0. To determine whether genes in the analyzed gene set are less likely to be SVGs than genes outside the set, we set M_0_: *a*_1_*_l_* ≥ 0 and M_1_: *a*_1_*_l_ <* 0. We quantify the evidence for enrichment using the Bayes factor (BF), defined as the ratio of posterior odds to prior odds:

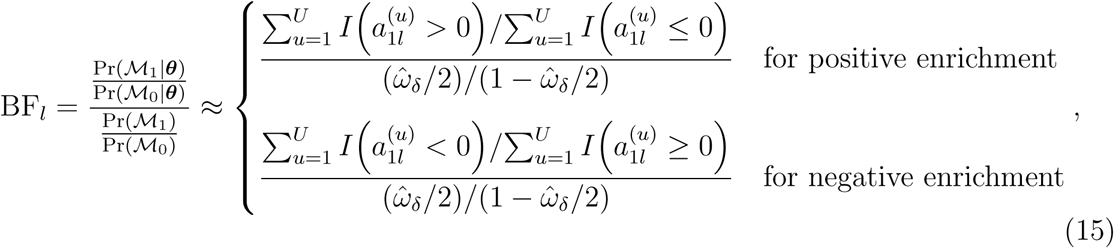

where {*a*_1*l*_^(*u*)^}*^U^_u_*_=1_ denotes the post-burn-in MCMC samples of *a*_1_*_l_* and *U* is the total number of retained iterations, and 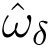 can be approximated by 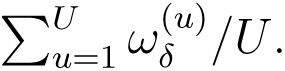 Larger values of BF_pos_ or BF_neg_ indicate stronger evidence for positive or negative enrichment, respectively, while accounting for uncertainty in latent SVG indicators and all model parameters. In this paper, our primary interest is in BF_pos_, since the main goal is to identify gene sets whose member genes are more likely to exhibit spatial variation than genes outside the set.

To aid interpretation, we adopt a reference BF threshold and report a gene set as significantly enriched when BF*_l_ >* 2.46, a commonly used reference that is comparable to the conventional threshold of *p* ≤ 0.05 [Sellke, 2012].

## 4 Simulation

To evaluate the performance of spaGSE, we first conducted a comprehensive simulation study designed to reflect realistic GSEA settings. With simulated summary statistics, we compared spaGSE with three widely used GSE methods including fGSEA [Korotkevich et al., 2016], clusterProfiler [Wu et al., 2021], and GOAT [Koopmans, 2024]. Across a wide range of simulation scenarios, spaGSE consistently achieved higher statistical power while maintaining strong false positive control compared to competing approaches.

We generated gene-level summary statistics under the Bayesian hierarchical model described in Section 2. Each simulated dataset contained *J* = 10, 000 genes. The baseline parameter *a*_0_*_l_* was chosen to reflect a sparse background level of spatial variation. We set *a*_0_*_l_* = −2, corresponding to an expected proportion of approximately 12% of genes outside of the analyzed gene set being SVGs. We further assessed the sensitivity of spaGSE to this baseline parameter by considering *a*_0_*_l_* ∈ {−0.5, −1, −3, −4}, corresponding to expected proportions of SVGs outside of the analyzed gene set of approximately 37.8%, 26.9%, 4.7%, and 1.8%, respectively. As reported in the Supplementary Section S4, spaGSE remained robust across these settings, maintaining high detection power of 98%–99% and well-controlled FPR, with the maximum observed FPR equal to 0.22%. The enrichment effect was generated to create either null gene sets, for which *a*_1_*_l_* = 0, or enriched gene sets, for which *a*_1_*_l_* /= 0. To evaluate performance under different levels of signal variability and gene set coverage, we considered two values of the gene-level variance parameter, *σ_θ_* ∈ {0.2, 0.5}, and three coverage rates (CRs), CR ∈ {0.3, 0.5, 0.7}. Here, the coverage rate represents the proportion of SVGs in the analyzed gene set and controls how strongly the pathway is represented among SVGs. We further fixed *σ_a_*= 10 to control the variability of gene set enrichment effects and set *ω_δ_* = 0.5 to generate the simulated datasets, indicating that each gene set has a 50% probability of nonzero enrichment effect.

Each simulation setting consisted of 1, 000 replicates, including 100 replicates with enriched gene sets (i.e., *a*_1_*_l_* /= 0) and 900 null replicates without enrichment (i.e., *a*_1_*_l_* = 0). This design allowed us to evaluate both statistical power under non-null scenarios and false positive control under null scenarios. Statistical power was defined as the proportion of truly enriched gene sets correctly identified, while false positive rate (FPR) was evaluated empirically as the proportion of null replicates incorrectly identified as enriched.

We implemented spaGSE *via* the Rcpp package. For prior specification, we assigned *ω_δ_* ∼ Beta(*a_ω_, b_ω_*) with *a_ω_* = 0.4 and *b_ω_* = 1.6, reflecting the prior belief that approximately 20% of gene sets are enriched. Flat priors were imposed on *a*_0_*_l_* and *a*_1_*_l_* by setting *σ*_0_ = 10 and *σ_a_* = 10. For posterior computation, the MH proposal variance for updating *a*_0_*_l_* was set to *τ*^(1)^*_a_*_0*l*_= 0.5. In the *add-delete* step for updating *a*_1_*_l_*, the proposal variance was set = 2, while conditional updates of *a*_1_*_l_* given *δ_l_* = 1 used a proposal variance of = 1. The MCMC algorithm was run for a total of 10, 000 iterations, with the first 5, 000 iterations discarded as burn-in.

To assess the convergence of the posterior sampler, we examined trace plots for the regression coefficients *a*_1_*_l_* and *a*_0_*_l_*. Across different simulation settings, the chains stabilized after burn-in and fluctuated around constant levels without apparent trends or long-term drifts (see Supplementary Figure S2). In particular, the trace plots for *a*_1_*_l_* showed adequate mixing around its posterior mean, while the trace plot for *a*_0_*_l_* remained centered near the true value. These diagnostics indicate that the proposed MCMC algorithm provides stable posterior sampling across the simulation settings considered.

For benchmarking, we compared spaGSE with fGSEA, GOAT, and clusterProfiler. For fGSEA and GOAT, the simulated gene-level *z*-scores were first converted to two-sided *p*-values according to *p_j_* = 2{1 − Φ(|*θ_j_*|)} and then used as inputs. fGSEA was implemented with 1, 000 permutations, and GOAT was applied using Bonferroni-adjusted *p*-values with a significance threshold of 0.05. For clusterProfiler, the simulated gene-level *z*-scores were analyzed through its GSEA interface.

The power comparison is shown in Figure 3. spaGSE consistently achieved the highest detection power across all six simulation settings. Specifically, the power of spaGSE remained stable at 97%–99% across different values of *σ_θ_* and CR. In contrast, fGSEA, clusterProfiler, and GOAT demonstrated moderate power of approximately 72%–73%. These results indicate that spaGSE improves power by approximately 36% relative to competing methods. Importantly, this gain was observed across different levels of signal variability and pathway coverage, demonstrating the strong robustness of spaGSE. We note that under the setting with CR = 70%, GOAT failed to produce a valid result, and hence no bar is displayed in Figure 3. By contrast, spaGSE produced stable results across all coverage levels.

**Figure 3:**
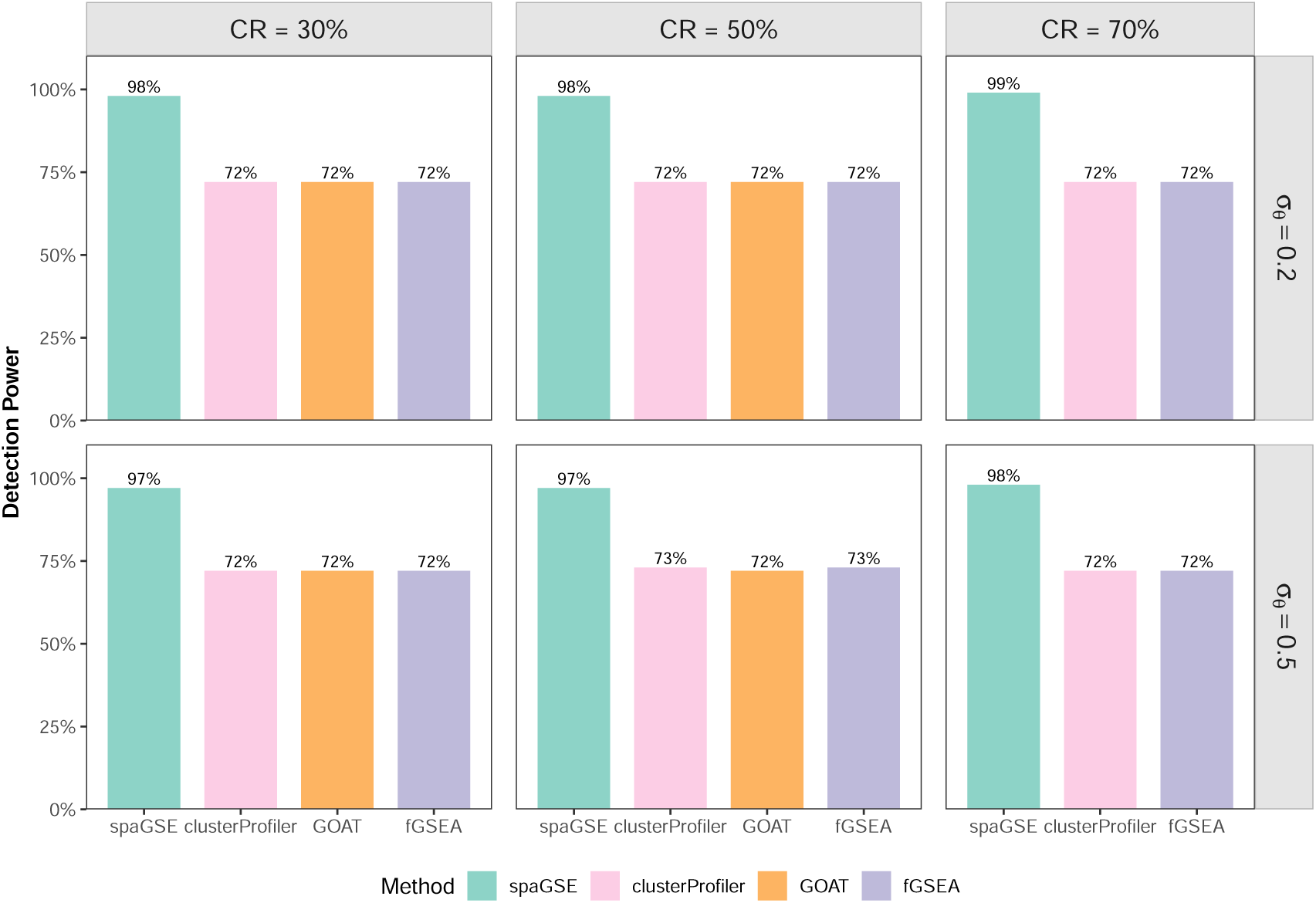
spaGSE is more powerful for GSE analysis than existing approaches in power simulations.

The false positive rate (FPR) comparison is shown in Figure 4. spaGSE maintained very low FPR across all settings, with values ranging from 0.00% to 0.22%. In comparison, the competing methods exhibited higher FPRs. The FPR of clusterProfiler ranged from 4.11% to 5.89%, GOAT ranged from 4.22% to 5.22% in settings where it returned valid results, and fGSEA remained between 4.11% and 6.44%. Moreover, the competing approaches showed a mild increase in FPR as the gene set coverage rate increased, while spaGSE remained nearly unchanged. Taken together, these results show that spaGSE is more powerful in identifying truly enriched spatial gene sets while providing much stronger control of false positives than existing alternatives.

**Figure 4:**
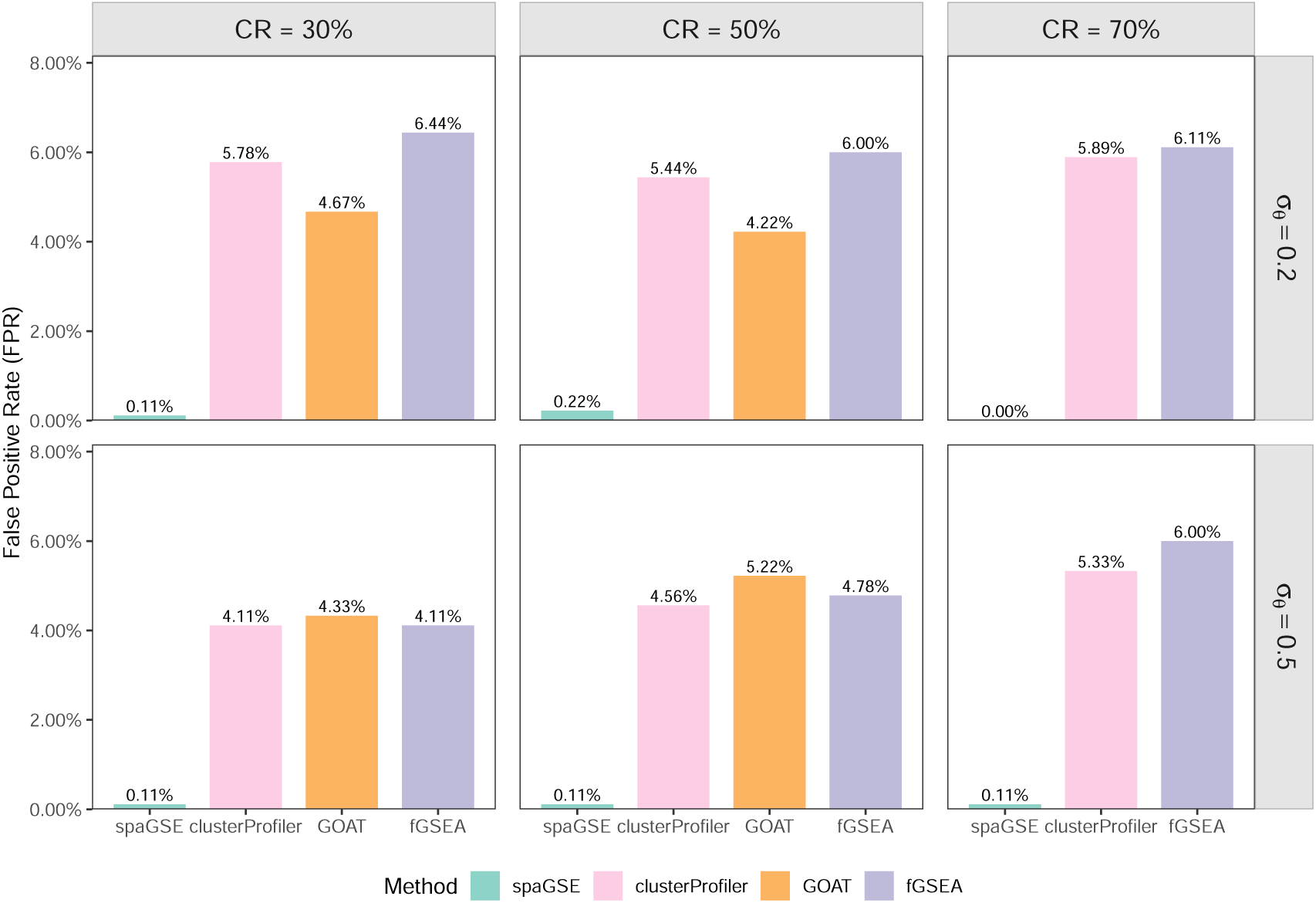
spaGSE achieves lower FPR for GSE analysis than existing approaches in FPR simulations.

We further evaluated the ability of spaGSE to detect true SVGs across simulation settings. Specifically, the SVG detection power was calculated as the proportion of true SVGs correctly identified among 100 non-null replicates (*a*_1_*_l_* /= 0). As shown in Supplementary

Figure S3, spaGSE achieved consistently high detection power when *σ_θ_* = 0.2, with power concentrated around 98% across different coverage rates. When *σ_θ_* = 0.5, the median power decreased to approximately 75%–80%, with moderately increased variability across replicates. This pattern suggests that gene-level SVG recovery becomes more challenging when the spatial summary statistics are noisier, although spaGSE still maintained moderate to high detection power across all coverage rates. These results indicate that spaGSE not only identifies enriched pathways with high power but also provides reliable inference on latent gene-level spatial status.

## 5 Applications

We applied spaGSE to four publicly available SRT datasets, including three generated using the 10x Genomics Visium platform and one generated using high-resolution Stereo-seq technology. For each dataset, raw count matrices were preprocessed using the Seurat package [Hao et al., 2021]. To reduce low-quality features and sparse signals, genes expressed in fewer than three cells and cells containing fewer than 100 detected genes were removed. We then applied SPARK-X to the filtered expression data to obtain gene-level spatial summary statistics. SPARK-X was used instead of SPARK because it provides improved computational efficiency for large-scale SRT datasets. Finally, spaGSE was fitted by integrating these gene-level summary statistics with predefined gene set annotations that are of biological interest.

### 5.1 Human PDAC 10x Visium Data

The first dataset we analyzed was a human PDAC SRT dataset [Moncada et al., 2020] containing expression measurements for 25, 753 genes across 428 spatial locations. After preprocessing, 16, 006 genes measured on 426 spots were retained for subsequent analysis. Using SPARK-X for spatial expression analysis, we identified 2, 625 SVGs, representing approximately 16.4% of all analyzed genes.

We then applied spaGSE to 485 cancer-related target gene sets obtained from the msigdbr database [Dolgalev et al., 2022]. Figure 5a summarizes the relationship between the gene set enrichment coefficient *a*_1_*_l_* and the corresponding BFs, log_10_(BF_pos_) and log_10_(BF_neg_). Gene sets identified as positively enriched by spaGSE generally have positive posterior estimates of *a*_1_*_l_* and large values of log_10_(BF_pos_), whereas gene sets with negative estimates tend to be supported by larger values of log_10_(BF_neg_). Most significant gene sets have posterior estimates of *a*_1_*_l_* between 2.0 and 4.0, corresponding to odds ratios of approximately 7.4 to 54.6 for genes within the gene set being spatially variable compared with those outside the gene set. These results indicate that the detected gene sets are strongly enriched for spatially structured transcriptional signals.

**Figure 5:**
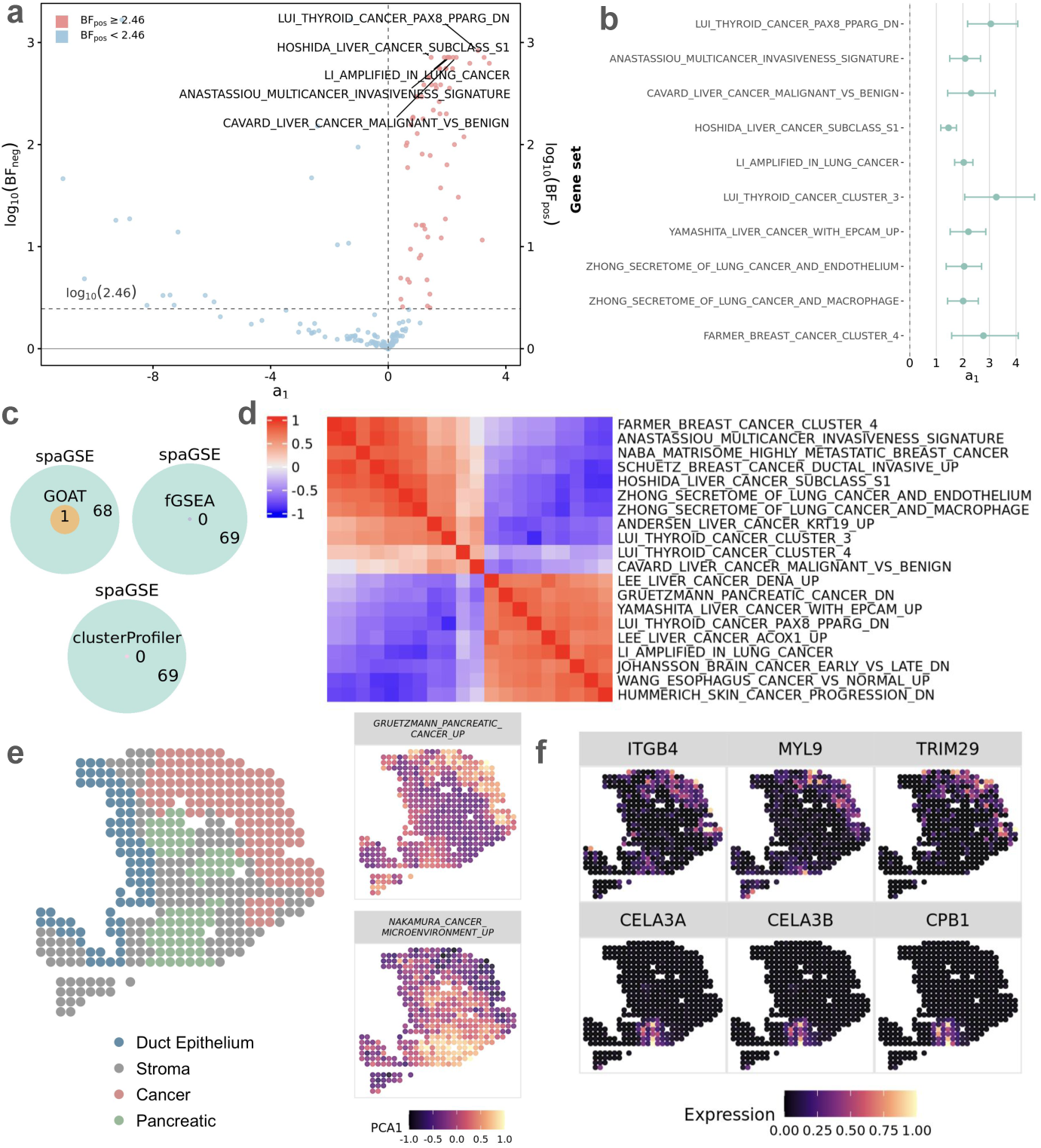
**Analysis results for the human PDAC data**: **a**. Volcano plot shows log_10_BF_pos_ for gene sets with *a*_1_*_l_ >* 0 and log_10_BF_neg_ for those with *a*_1_*_l_ <* 0 from spaGSE for GSE analysis. Gene sets colored by red are identified as statistically significant by spaGSE. **b**. Posterior means and 95% CIs of *a*_1_*_l_* for the top ten representative gene sets detected by spaGSE. **c**. Venn graphs show gene sets detected by spaGSE, GOAT, and fGSEA. **d**. A heatmap shows the correlation for selected top 20 gene sets based on the first principal component after PCA. Most gene sets are clustered into two groups. **e**. Spatial plots display gene set expression patterns. **f**. Spatial plots show three selected SVGs for each pathway presented in e.

To further assess the stability of the estimated enrichment effects, we examined the posterior means of *a*_1_*_l_* and their corresponding 95% CIs for the top ten representative gene sets (Figure 5b). Notably, most CIs are relatively narrow and clearly separated from zero, suggesting stable posterior inference for these gene sets. Consistent with the simulation results, spaGSE identified more significantly enriched gene sets than competing GSE methods (Figure 5c). For spaGSE, significance was assessed using a reference Bayes factor threshold of 2.46, which is commonly regarded as comparable to the conventional criterion of *p* ≤ 0.05. Under this criterion, spaGSE identified 69 significant gene sets. In contrast, using the corresponding *p*-value-based criteria, GOAT identified only one, and the other two methods failed to detect any significant gene sets. This comparison suggests that spaGSE can improve detection power in settings where pathway-level spatial signals are distributed across multiple genes but may not be sufficiently captured by conventional

### GSE methods

We next examined the relationships among the gene sets detected by spaGSE. For each significant gene set, we summarized its spatial activity profile using the first principal component score of the expression matrix restricted to genes in that set. We then computed pairwise Pearson correlations between these gene set activity profiles. The correlation heatmap of the top 20 representative gene sets reveals two major clusters (Figure 5d). The first cluster is enriched for invasion-and tumor microenvironment-related signatures, including ANASTASSIOU MULTICANCER INVASIVENESS SIGNATURE, and NABA MATRISOME HIGHLY METASTATIC BREAST CANCER. In contrast, the second cluster contains broader tumor-associated transcriptional programs, such as LEE LIVER CANCER ACOX1 UP, and YAMASHITA LIVER CANCER WITH EPCAM UP. These clusters indicate that spaGSE not only identifies individual enriched pathways but also groups spatially coherent cancer-related programs into biologically interpretable clusters. A complete correlation heatmap for all 69 gene sets identified by spaGSE is provided in the Supplementary Figure S6.

Several identified gene sets are biologically consistent with known features of PDAC heterogeneity. For example, the pancreatic cancer-related gene set GRUETZMANN PANCRE-ATIC CANCER UP and the microenvironment-related gene set NAKAMURA CANCER MICROENVIRONMENT UP exhibit distinct spatial gene set activity patterns (Figure 5e). Specifically, GRUETZMANN PANCREATIC CANCER UP shows strong activity in the tumor region, consistent with its reported association with PDAC [Grützmann et al., 2005]. In contrast, NAKAMURA CANCER MICROENVIRONMENT UP shows elevated activ-ity around both the tumor and pancreatic regions. This gene set was originally defined from genes that were upregulated when pancreatic cancer cells were grown as orthotopic tumors in the pancreas, compared with cells grown in vitro [Nakamura et al., 2007]. In the original study, this comparison was designed to evaluate how the local tissue microenvironment influences gene expression in pancreatic cancer cells. This spatial pattern is consistent with its reported association with microenvironment-induced transcriptional changes in pancreatic cancer cells.

The gene-level spatial expression patterns further support the gene set-level findings and the modeling motivation of spaGSE. Representative SVGs within the same enriched pathway exhibit coherent spatial expression patterns (Figure 5f). This indicates that the pathway-level enrichment detected by spaGSE is driven by coordinated spatial variation among multiple genes rather than by isolated individual signals. Additional SVGs from these two pathways are shown in Supplementary Figures S7 and S8, where their overall spatial coherence is illustrated more comprehensively. In summary, this case study demonstrates that spaGSE can identify interpretable gene sets whose member genes show coordinated spatial organization and are closely related to key features of PDAC tumor biology.

### 5.2 Human Breast Cancer 10x Visium Data

The second SRT dataset we analyzed was a human breast cancer 10x Visium dataset [Jiang et al., 2024], consisting of expression measurements for 17, 943 genes across 1, 272 spatial locations. After preprocessing, 15, 852 genes remained for subsequent analysis. We applied SPARK-X to the filtered expression matrix to obtain gene-level spatial association statistics and identified 9, 528 SVGs, representing 60.1% of all analyzed genes.

We then applied spaGSE to 126 breast cancer-related target gene sets from the msigdbr database. Figure 6a summarizes the relationship between the estimated gene set enrichment coefficients *a*_1_*_l_* and the corresponding BFs, log_10_(BF_pos_) and log_10_(BF_neg_). Most gene sets identified as significant by spaGSE have positive enrichment coefficients and are associated with relatively large log_10_(BF_pos_) values, while gene sets with negative *a*_1_*_l_* values tend to be supported by larger values of log_10_(BF_neg_). Most significant gene sets have posterior estimates of *a*_1_*_l_* between 1.0 and 3.0, corresponding to odds ratios of approximately 2.7 to 20.1 for genes within the gene set being spatially variable compared with those outside the gene sets. These results indicate that the detected gene sets are enriched for spatially organized transcriptional signals in the breast cancer tissue section. We also presented the posterior means of *a*_1_*_l_* and their corresponding 95% CIs for the top ten representative gene sets (Figure 6b). Notably, most CIs are relatively narrow and separated from zero, indicating stable posterior estimation. Consistent with the simulation results, spaGSE identified more significantly enriched gene sets than competing GSE methods (Figure 6c). Applying the same BF cutoff, spaGSE identified 45 significantly enriched gene sets, whereas GOAT and clusterProfiler identified only 33 and 25 gene sets, respectively. In contrast, fGSEA did not identify any target gene set as significant. This comparison suggests that spaGSE can detect pathway-level spatial signals that may be missed or less consistently recovered by conventional GSE methods.

**Figure 6:**
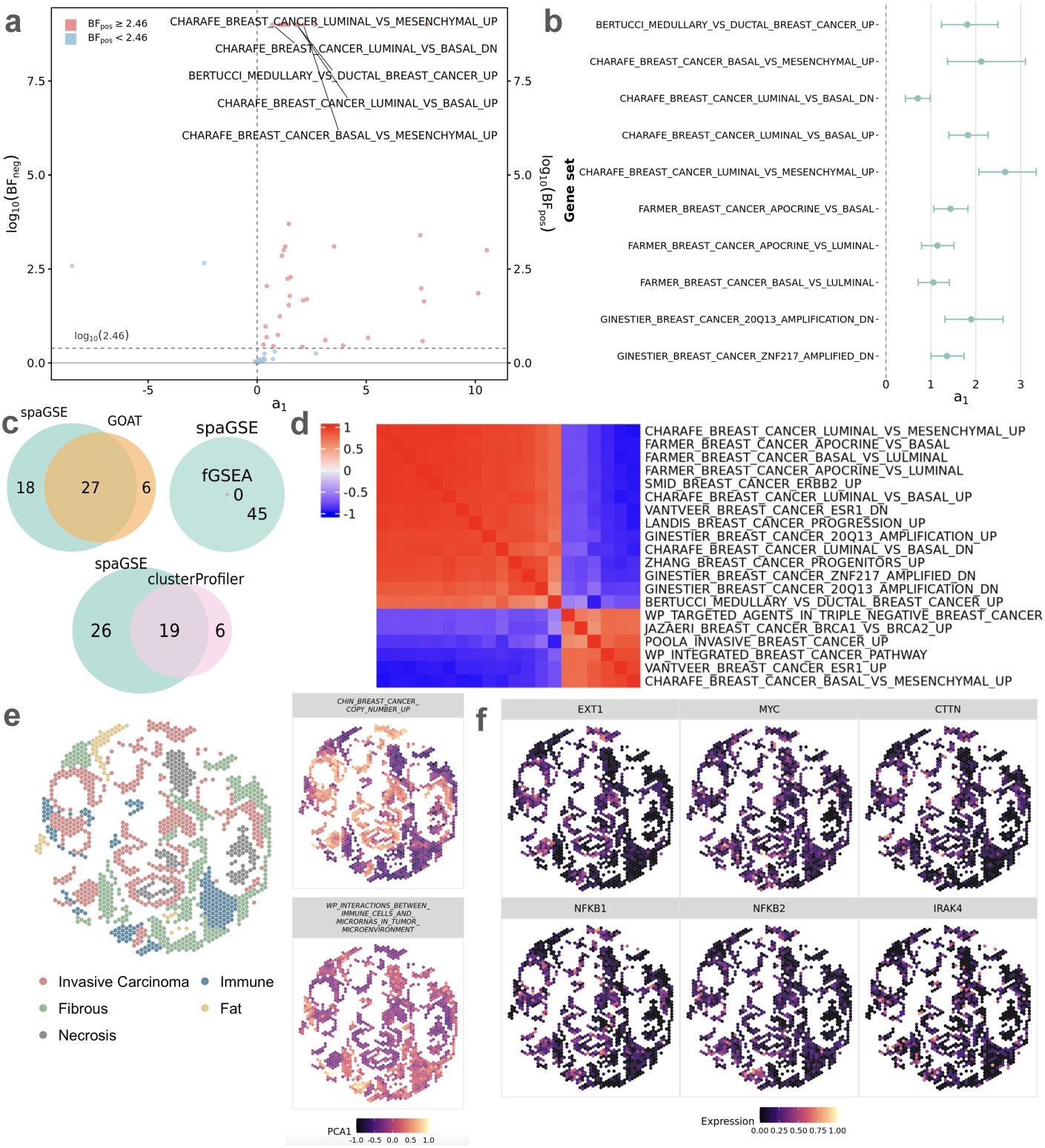
**Analysis results for the FFPE human breast cancer data**: **a**. Volcano plot shows log_10_BF_pos_ for gene sets with *a*_1_*_l_ >* 0 and log_10_BF_neg_ for those with *a*_1_*_l_ <* 0 from spaGSE for GSE analysis. Gene sets colored by red are identified as statistically significant by spaGSE. **b**. Posterior means and 95% CIs of *a*_1_*_l_* for the top ten representative gene sets detected by spaGSE.**c**. Venn graphs show gene sets detected by spaGSE, GOAT, and fGSEA. **d**. A heatmap shows the correlation for selected top 20 gene sets based on the first principal component after PCA. The gene sets are clearly clustered into two groups. **e**. Spatial plots display gene set expression patterns. **f**. Spatial plots show three selected SVGs for each pathway presented in e.

We further examined the relationships among the gene sets detected by spaGSE. The clustered correlation heatmap of the top 20 representative gene sets reveals two major pathway modules (Figure 6d). One module is primarily composed of luminal-, apocrine-, and ESR1-related breast cancer signatures, while the other was enriched for basal-like, invasive, and integrated breast cancer pathway signatures. This structured correlation pattern suggests that the enriched gene sets identified by spaGSE capture biologically coherent transcriptional programs. A complete correlation heatmap for all 45 significant gene sets identified by spaGSE is provided in the Supplementary Figure S9.

Several identified gene sets also provide biologically meaningful insights into spatial heterogeneity in the FFPE breast cancer tissue. For example, the breast cancer-related gene set CHIN BREAST CANCER COPY NUMBER UP and the immune-related gene set WP INTERACTIONS BETWEEN IMMUNE CELLS AND MICRORNAS IN TUMOR MICROENVIRONMENT exhibit clearly distinct spatial localization patterns (Figure 6e). Specifically, CHIN BREAST CANCER COPY NUMBER UP shows strong activity around the invasive carcinoma region, consistent with its close relevance to breast cancer biology, while WP INTERACTIONS BETWEEN IMMUNE CELLS AND MICRORNAS IN TUMOR MICROENVIRONMENT displays stronger activity in the immune cell-enriched region, consistent with the role of immune regulation and microRNA-mediated interactions in the tumor microenvironment (Chou et al. [2013]). These spatial patterns suggest that spaGSE can identify pathway-level signals corresponding to distinct tumor and immune components within the same tissue section. The gene-level spatial expression patterns further support the pathway-level findings. Representative SVGs within the same enriched pathway exhibit similar spatial expression patterns (Figure 6f). Additional SVGs with clear spatial signals from these two pathways are shown in the Supplementary Figures S10 and S11, where the overall spatial coherence is illustrated more comprehensively. In summary, this case study in a different tumor setting shows that spaGSE can identify biologically interpretable pathways that are supported by coordinated spatial expression patterns and closely aligned with key features of breast cancer biology.

### 5.3 Human Prostate Cancer 10x Visium Data

The third SRT dataset we analyzed was a human prostate cancer 10x Visium dataset [Jiang et al., 2024], consisting of gene expression measurements for 17, 943 genes across 4, 371 spatial locations. Since the primary goal of this analysis was to examine pathway-level spatial heterogeneity within malignant tissue, we restricted the analysis to the annotated tumor region, which contained 1, 659 spatial locations. After preprocessing, 15, 165 genes were remained for downstream analysis. Applying SPARK-X to the filtered expression matrix identified 5, 431 SVGs, corresponding to 35.8% of all analyzed genes.

We applied spaGSE to 485 cancer-related target gene sets obtained from the msigdbr database. Figure 7a shows the association between the estimated gene set enrichment co-efficient *a*_1_*_l_* and the BFs, log_10_(BF_pos_)(log_10_(BF_neg_)). As expected, gene sets identified as significant by spaGSE have positive enrichment coefficients and are associated with relatively large log_10_(BF_pos_) values, while gene sets with negative *a*_1_*_l_* estimates are supported by larger values of log_10_(BF_neg_). Among the significant gene sets, most posterior estimates of *a*_1_*_l_* fall between 1.0 and 4.0, corresponding to odds ratios of approximately 2.7 to 54.6 for genes within the gene set being spatially variable compared with those outside the set. This range indicates that many detected pathways show substantial enrichment for spatially structured transcriptional variation within the tumor region.

**Figure 7:**
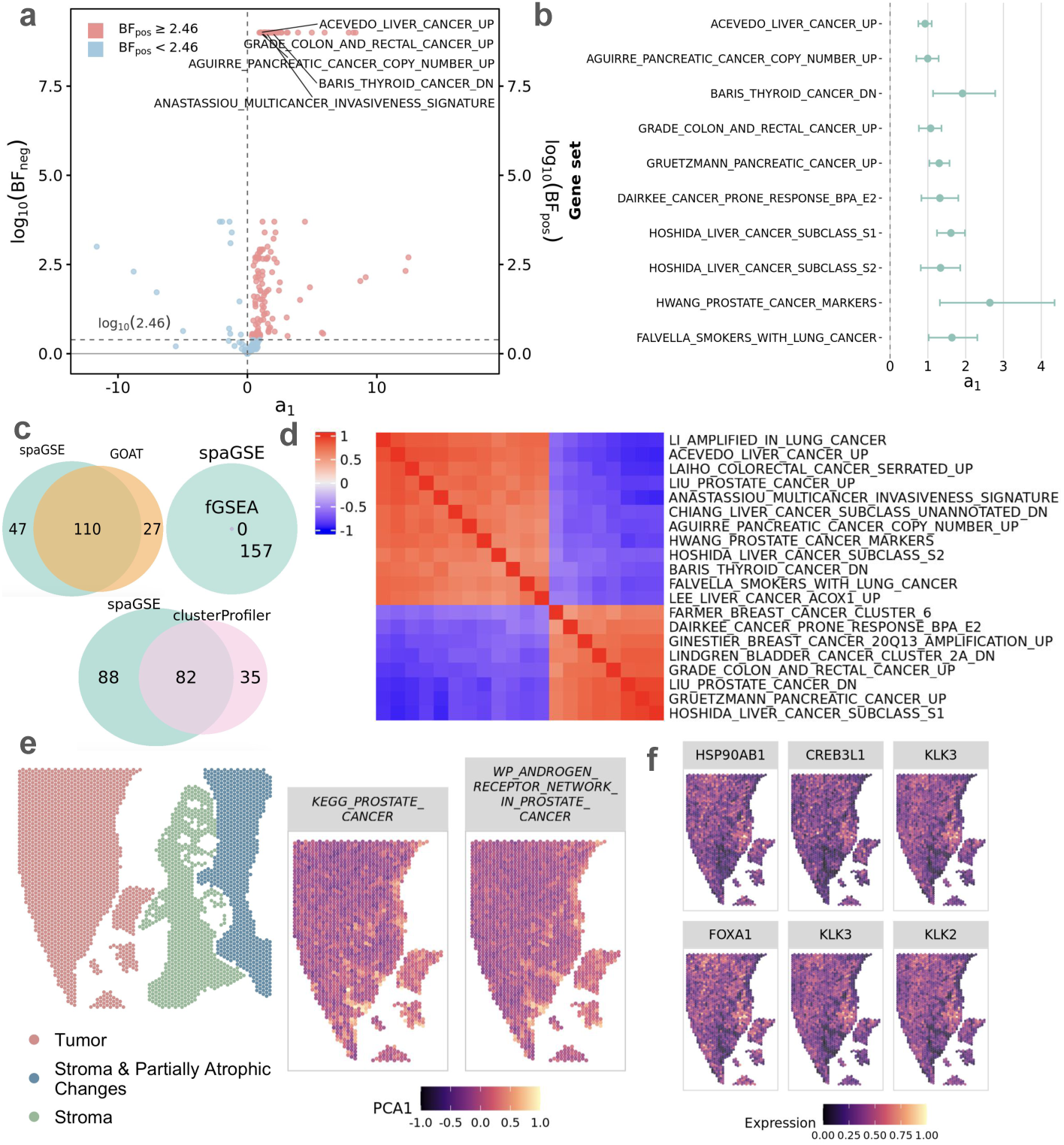
**Analysis results for the FFPE human prostate cancer data**: **a**. Volcano plot shows log_10_BF_pos_ for gene sets with *a*_1_*_l_ >* 0 and log_10_BF_neg_ for those with *a*_1_*_l_ <* 0 from spaGSE for GSE analysis. Gene sets colored by red are identified as statistically significant by spaGSE. **b**. Posterior means and 95% CIs of *a*_1_*_l_* for the top ten representative gene sets detected by spaGSE.**c**. Venn graphs show gene sets detected by spaGSE, GOAT, and fGSEA. **d**. A heatmap shows the correlation for selected top 20 gene sets based on the first principal component after PCA. The gene sets are clearly clustered into two groups. **e**. Spatial plots display gene set expression patterns. **f**. Spatial plots show three selected SVGs for each pathway presented in e.

The posterior means of *a*_1_*_l_* for the top ten representative gene sets are shown in Figure 7b, together with their corresponding 95% CIs. Notably, most CIs are relatively narrow and away from zero, indicating stable enrichment estimates for these pathways. In comparison with existing GSE methods, spaGSE identified a larger number of significant gene sets (Figure 7c). Applying the same BF cutoff, spaGSE identified 157 significant gene sets, whereas GOAT and clusterProfiler identified only 137 and 117, respectively. In contrast, fGSEA did not identify any target gene set as significant. Notably, 14 of the gene sets identified by spaGSE were directly related to prostate cancer, supporting the biological relevance of the detected pathway-level spatial signals.

Among the top 20 representative gene sets identified by spaGSE, the clustered correlation heatmap reveals two major groups of pathways (Figure 7d). One group is enriched for prostate cancer-related, invasiveness-related, and cancer marker signatures, whereas the other is composed primarily of non-prostate cancer signatures derived from other tumor types but reflecting shared malignant processes. This pattern suggests that the enriched pathways capture both disease-specific and more general cancer-associated transcriptional programs with spatial structure. The complete correlation heatmap for all 157 significant gene sets is provided in the Supplementary Figure S12.

Several of the identified gene sets also provide biologically meaningful insights into the spatial organization of the prostate tumor region. In particular, two prostate cancer-related gene sets KEGG PROSTATE CANCER and WP ANDROGEN RECEPTOR NETWORK IN PROSTATE CANCER show strong gene set activity around the tumor region and exhibit similar spatial localization patterns (Figure 7e), consistent with their close relevance to prostate cancer biology [Fan et al., 2018]. These pathway-level signals are further supported by the gene-level results, where representative SVGs from the same gene sets display similar spatial expression patterns (Figure 7f). Additional SVGs from these two pathways are shown in the Supplementary Figures S13 and S14, further supporting the coordinated spatial variation of member genes within each pathway. Together, these results show that spaGSE identifies biologically meaningful pathways with coordinated spatial expression patterns and highlights key transcriptional programs underlying the spatial organization of prostate cancer tissue.

### 5.4 Mouse Embryo Stereo-seq Data

Finally, we analyzed a mouse embryo Stereo-seq dataset at developmental stage E16.5 [Chen et al., 2022]. Compared with the Visium datasets, this dataset provides a substantially larger and more spatially detailed transcriptomic map, with 23, 761 genes measured across 121, 767 spatial locations, thereby posing a greater computational challenge. After preprocessing, 23, 685 genes were retained for subsequent analysis. SPARK-X identified 18,844 SVGs, corresponding to 79.6% of the analyzed genes, reflecting the extensive spatial organization of gene expression during embryonic development.

We then applied spaGSE to 311 curated target gene sets related to cardiac and muscle development, neural differentiation, skeletal and cartilage formation, and vascular organization collected from the msigdbr database. Figure 8a shows the posterior enrichment estimates *a*_1_*_l_* together with the corresponding BFs. Most gene sets identified as significant by spaGSE have positive enrichment coefficients and are associated with relatively large log_10_(BF_pos_) values, while only a few gene sets with negative *a*_1_*_l_* values show stronger evidence through log_10_(BF_neg_). Moreover, many significant gene sets have large estimated *a*_1_*_l_* values between 8.0 and 12.0, suggesting strong positive enrichment of SVGs within these pathways. We also presented the posterior means of *a*_1_*_l_* and their corresponding 95% CIs for the top ten representative gene sets (Figure 8b). Notably, most CIs are relatively narrow, indicating stable posterior estimation. Consistent with the simulation results, spaGSE again identified more significantly enriched target gene sets than the other GSE methods (Figure 8c). Applying the same BF cutoff, spaGSE identified 272 significantly enriched gene sets, whereas GOAT, clusterProfiler, and fGSEA identified only 136, 170, and 246, respectively.

**Figure 8:**
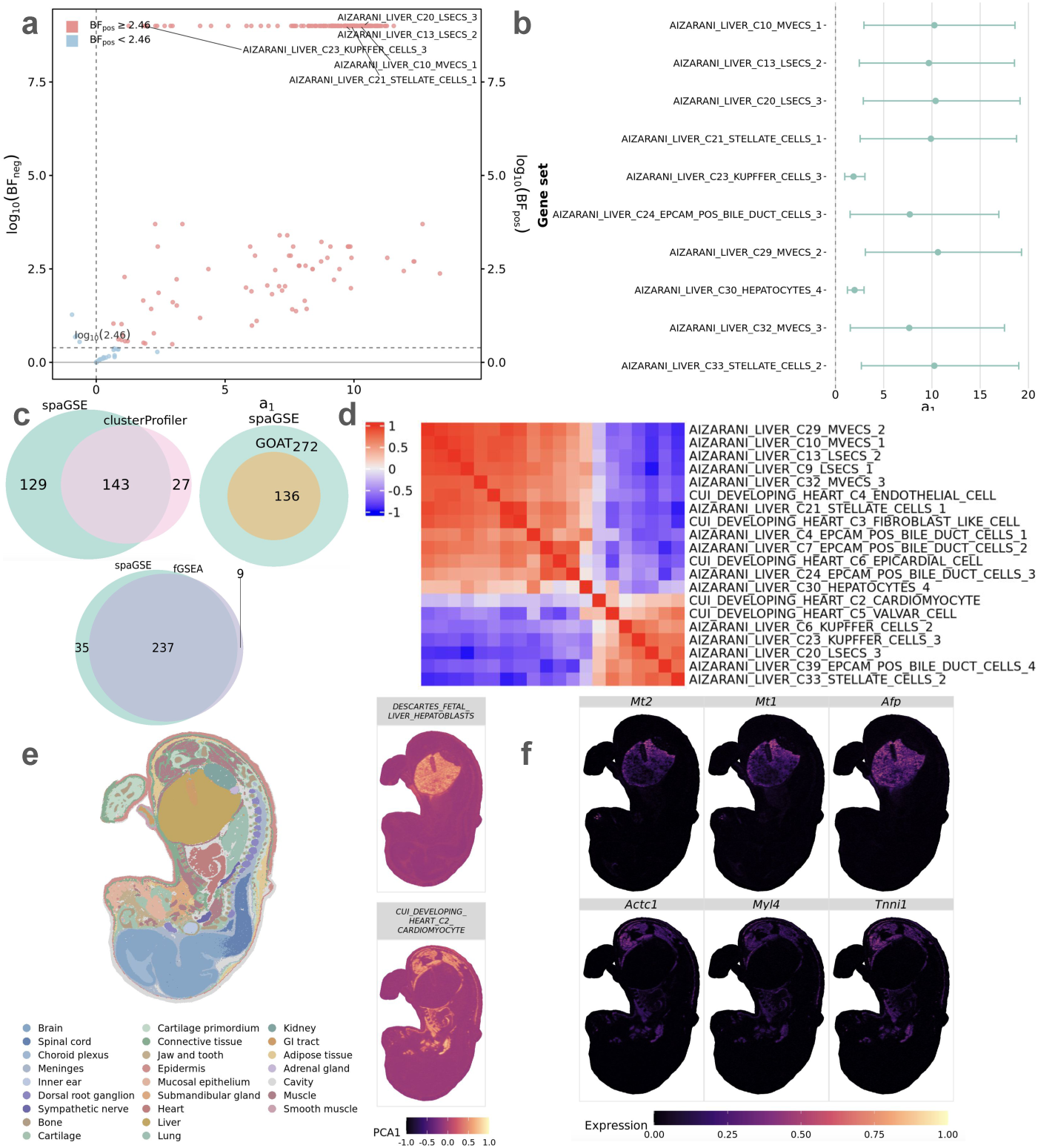
A**n**alysis **results for the mouse embryo data at stage E16.5**: **a**. Volcano plot shows log_10_BF_pos_ for gene sets with *a*_1_*_l_ >* 0 and log_10_BF_neg_ for those with *a*_1_*_l_ <* 0 from spaGSE for GSE analysis. Gene sets colored by red are identified as statistically significant by spaGSE. **b**. Posterior means and 95% CIs of *a*_1_*_l_* for the top ten representative gene sets detected by spaGSE.**c**. Venn graphs show gene sets detected by spaGSE, GOAT, and fGSEA. **d**. A heatmap shows the correlation for selected top 20 gene sets based on the first principal component after PCA. The gene sets are clearly clustered into two groups. **e**. Spatial plots display gene set expression patterns. **f**. Spatial plots show three selected SVGs for each pathway presented in e.

The top 20 representative gene sets identified by spaGSE show a clear modular structure in their spatial gene set activity profiles (Figure 8d). The clustered correlation heatmap separates these gene sets into two major groups. One group is dominated by vascular, stromal, and epithelial-associated programs, whereas the other is enriched for parenchymal, immune, and contractile programs. This organization suggests that the significant gene sets reflect distinct but related developmental processes across the E16.5 embryo. The complete correlation heatmap for all 272 significant gene sets identified by spaGSE is provided in the Supplementary Figure S15.

Several identified gene sets also provide biologically meaningful insights into spatial developmental heterogeneity. In particular, gene sets DESCARTES FETAL LIVER HEPATO-BLASTS and CUI DEVELOPING HEART C2 CARDIOMYOCYTE exhibit clearly distinct spatial patterns (Figure 8e). Specifically, DESCARTES FETAL LIVER HEPATOBLASTS shows highly localized expression in the annotated liver region, consistent with its connection to liver development. In contrast, CUI DEVELOPING HEART C2 CARDIOMYOCYTE shows strong activity around the annotated heart region and nearby muscle-related structures, consistent with its relevance to heart-related development [Raso et al., 2021, Yanget al., 2023]. At the gene level, representative SVGs from these pathways exhibit similar spatial expression patterns (Figure 8f), indicating that the pathway-level signals are supported by coordinated spatial variation among multiple member genes. The full sets of additional SVGs with clear spatial signals from these two pathways are provided in the Supplementary Figures S16 and S17. In summary, these results show that spaGSE identifies biologically meaningful pathways that are supported by coordinated spatial expression patterns and closely aligned with key features of embryonic tissue development.

## 6 Conclusion

In this paper, we developed spaGSE, a Bayesian hierarchical model for GSEA in SRT. By modeling gene-level summary statistics derived from spatial expression analysis, spaGSE links spatial signals in SRT data with a predefined gene set database and enables pathway-level inference on spatial enrichment. Specifically, spaGSE uses a logistic regression formulation to quantify the association between genelevel spatial signals and gene set membership, and imposes a spike-and-slab prior on the enrichment coefficient to support robust and interpretable inference. Through comprehensive simulation studies and analyses of four real SRT datasets from different tissue types, we demonstrated that spaGSE achieved superior performance compared to existing methods, with higher power and better control of false positives. In realdata applications, spaGSE identified biologically meaningful gene sets with coherent spatial organization that were missed by competing approaches, including tumor-associated, immune-related, and developmental pathways. These results demonstrate that integrating spatial information and jointly modeling gene-level summary statistics into GSE analysis can improve both statistical performance and biological interpretability, making spaGSE a useful tool for studying pathway-level spatial variation in SRT studies.

There are several important future extensions for spaGSE. First, the current implementation of spaGSE takes gene-level summary statistics from SPARK as input. Although this design is computationally efficient, the performance of spaGSE may still be influenced by the quality of the upstream spatial expression analysis. Therefore, future work could extend the framework to accommodate summary statistics from different SVG detection methods or alternative measures of gene-level spatial evidence, thereby improving its robustness and broadening its applicability. Second, spaGSE could be extended to incorporate external spatial or multimodal information, such as histology images, and cell-type annotations, to improve the accuracy and biological relevance of spatial pathway enrichment analysis. Leveraging these complementary data sources may provide a more comprehensive understanding of tissue organization and cellular microenvironments. Finally, an important future direction is to generalize spaGSE to multi-sample or multi-group settings. Many SRT studies now include multiple patients, tissue sections, disease states, or treatment conditions. Extending spaGSE to model shared and condition-specific pathway-level spatial enrichment would enable systematic comparisons across biological groups and support the study of spatial pathway heterogeneity at the cohort level. These extensions would further broaden the applicability of spaGSE and strengthen its utility for studying spatially organized biological processes in complex tissues.

## Data Availability Statement

The authors analyzed four publicly available SRT data. Raw count matrices and spatial information for four SRT data from 10x Visium and Stereo-seq are accessible at https://github.com/YMa-lab/spaGSE/tree/main/application. An open-source implementation of the spaGSE algorithm in R/C++ is also available at https://github.com/YMa-lab/spaGSE/tree/main.

## Author contributions

Y.M. conceived the idea and provided funding support. Z.X. and Y.G. developed the method. Z.X. implemented the software, performed simulations, and analyzed real data. Z.X., Y.M., and Q.L. wrote the paper.

## References

Sikta Das Adhikari et al. Recent advances in spatially variable gene detection in spatial transcriptomics. Computational and Structural Biotechnology Journal, 23:883–891, 2024.

Yuzhou Chang et al. Graph Fourier transform for spatial omics representation and analyses of complex organs. Nature Communications, 15(1):7467, 2024.

Ao Chen et al. Spatiotemporal transcriptomic atlas of mouse organogenesis using DNA nanoball-patterned arrays. Cell, 185(10):1777–1792, 2022.

Yanguang Chen. Spatial autocorrelation equation based on Moran’s index. Scientific reports, 13 (1):19296, 2023.

Jonathan Chou, Payam Shahi, and Zena Werb. microrna-mediated regulation of the tumor microenvironment. Cell cycle, 12(20):3262–3271, 2013.

David DeTomaso and Nir Yosef. Hotspot identifies informative gene modules across modalities of single-cell genomics. Cell systems, 12(5):446–456, 2021.

Igor Dolgalev et al. msigdbr: MSigDB gene sets for multiple organisms in a tidy data format. R package version, 7(1):532–2, 2022.

Daniel Edsgärd, et al. Identification of spatial expression trends in single-cell gene expression data. Nature methods, 15(5):339–342, 2018.

Shutong Fan et al. Identification of the key genes and pathways in prostate cancer. Oncology Letters, 16(5):6663–6669, 2018.

Robert Grützmann, et al. Meta-analysis of microarray data on pancreatic cancer defines a set of commonly dysregulated genes. Oncogene, 24(32):5079–5088, 2005.

Yuhan Hao et al. Integrated analysis of multimodal single-cell data. Cell, 184(13):3573–3587, 2021.

Guichen Huang et al. The mechanism of ITGB4 in tumor migration and invasion. Frontiers in Oncology, 14:1421902, 2024.

Sanjay Jain and Michael T Eadon. Spatial transcriptomics in health and disease. Nature reviews nephrology, 20(10):659–671, 2024.

Xi Jiang et al. A Bayesian modified ising model for identifying spatially variable genes from spatial transcriptomics data. Statistics in medicine, 41(23):4647–4665, 2022.

Xi Jiang et al. iIMPACT: Integrating image and molecular profiles for spatial transcriptomics analysis, May 2024. URL 10.5281/zenodo.11117768.

Frank Koopmans. GOAT: efficient and robust identification of gene set enrichment. Communications biology, 7(1):744, 2024.

Gennady Korotkevich et al. Fast gene set enrichment analysis. biorxiv, page 060012, 2016.

Qiongyu Li et al. Spatial transcriptomics for tumor heterogeneity analysis. Frontiers in Genetics, 13:906158, 2022.

Qiwei Li, Minzhe Zhang, Yang Xie, and Guanghua Xiao. Bayesian modeling of spatial molecular profiling data via Gaussian process. Bioinformatics, 37(22):4129–4136, 2021.

Zhijian Li et al. Systematic benchmarking of computational methods to identify spatially variable genes. Genome Biology, 26(1):285, 2025.

Long Liu et al. TRIM29 promotes pancreatic cancer MVI via I*κ*B*α* k48-ubiquitination and NF-*κ*B activation in CXCL5^+^ epithelial cells. Molecular Cancer, 2026.

Minghe Lv et al. The landscape of prognostic and immunological role of myosin light chain 9 (MYL9) in human tumors. Immunity, Inflammation and Disease, 10(2):241–254, 2022.

Ying Ma et al. Integrative differential expression and gene set enrichment analysis using summary statistics for scRNA-seq studies. Nature communications, 11(1):1585, 2020.

Jeffrey R Moffitt, Emma Lundberg, and Holger Heyn. The emerging landscape of spatial profiling technologies. Nature Reviews Genetics, 23(12):741–759, 2022.

Reuben Moncada et al. Integrating microarray-based spatial transcriptomics and single-cell RNA-seq reveals tissue architecture in pancreatic ductal adenocarcinomas. Nature biotechnology, 38 (3):333–342, 2020.

Lambda Moses and Lior Pachter. Museum of spatial transcriptomics. Nature methods, 19(5): 534–546, 2022.

Toru Nakamura et al. Gene expression profile of metastatic human pancreatic cancer cells depends on the organ microenvironment. Cancer research, 67(1):139–148, 2007.

Andrea Raso et al. A microRNA program regulates the balance between cardiomyocyte hyperplasia and hypertrophy and stimulates cardiac regeneration. Nature communications, 12(1): 4808, 2021.

Thomas M Sellke. On the interpretation of p-values. Technical report, Tech. Rep. Department of Statistics, Purdue University, 2012.

Aravind Subramanian et al. Gene set enrichment analysis: a knowledge-based approach for interpreting genome-wide expression profiles. Proceedings of the national academy of sciences, 102(43):15545–15550, 2005.

Shiquan Sun et al. Statistical analysis of spatial expression patterns for spatially resolved transcriptomic studies. Nature methods, 17(2):193–200, 2020.

Valentine Svensson et al. SpatialDE: identification of spatially variable genes. Nature methods, 15(5):343–346, 2018.

Tianzhi Wu et al. clusterprofiler 4.0: A universal enrichment tool for interpreting omics data. The innovation, 2(3), 2021.

Guanao Yan et al. Categorization of 34 computational methods to detect spatially variable genes from spatially resolved transcriptomics data. Nature Communications, 16(1):1141, 2025.

Li Yang et al. The default and directed pathways of hepatoblast differentiation involve distinct epigenomic mechanisms. Developmental Cell, 58(18):1688–1700, 2023.

Xiang Zhou, Peter Carbonetto, and Matthew Stephens. Polygenic modeling with Bayesian sparse linear mixed models. PLoS genetics, 9(2):e1003264, 2013.

Jiaqiang Zhu et al. SPARK-X: non-parametric modeling enables scalable and robust detection of spatial expression patterns for large spatial transcriptomic studies. Genome biology, 22(1):184, 2021.

