## Supplementary material for "Spatial Gene Set Enrichment Analysis with Applications to Spatially Resolved Transcriptomic Data": NA

### Supplementary Materials for "Spatial Gene Set Enrichment Analysis with Applications to Spatially Resolved Transcriptomics Data"

Zizhao Xie[1], Yanghong Guo[2], Qiwei Li[2*], Ying Ma[1,3*]
[1]Department of Biostatistics, Brown University
[2]Department of Mathematical Sciences, The University of Texas at Dallas
[3]Center for Computational Molecular Biology, Brown University

#### S1 Notation Table

Supplementary Table S1 summarizes the main notation used in the spaGSE model. The table includes notation for the input data, latent SVG model, gene set enrichment model, prior parameters, MCMC algorithm, and posterior inference quantities.

#### S2 Spatial Autocorrelation of Pathway-Member SVGs

Figure S1 compares gene-level Moran's I values between SVGs within the GRUETZMANN_PANCREATIC_CANCER_UP gene set and SVGs outside this gene set. SVGs belonging to this pathway showed substantially higher spatial autocorrelation, with a median Moran's I around 0.30, whereas SVGs outside the pathway were concentrated near lower Moran's $I$ values aroud 0.05. This pattern suggests that spatially variable genes within the PDAC-related pathway exhibit more coherent spatial organization than the broader background of SVGs. These results provide additional support for the motivation of spaGSE, which leverages gene-set membership to identify pathways whose member genes share stronger spatial structure.

#### S3 MCMC Trace Plots

This section provides trace plots for representative model parameters under the prior specification and MCMC algorithm described in Section 4.2 of the main text. These plots are used to assess mixing behavior and convergence of the posterior sampling procedure.

Figure S2 presents the MCMC trace plots of one replicate for the gene set enrichment coefficients $a_{0l}$ and $a_{1l}$ under the setting $\sigma_\theta = 0.2, \mathrm{CR} = 30\%$. In panel A, the Markov chain for $a_{1l}$ rapidly reaches a stable region around the posterior mean 4.92 after a short initial adjustment and then fluctuates narrowly around this level throughout the remaining iterations. In panel B, the chain for $a_{0l}$ similarly stabilizes quickly around -2.0 and shows no apparent long-term drift. The dashed horizontal lines indicate the posterior sample means of the two parameters. Overall, both trace plots exhibit stable behavior, good mixing, and no visible trend after burn-in, suggesting well convergence of the MCMC algorithm under this simulation setting.

#### S4 SVG detection power across simulation settings

We further evaluated the SVG detection performance of spaGSE at the gene level. For each gene $j$, the posterior inclusion probability (PIP) was defined as the posterior probability that the gene is spatially variable, that is,

$$\text{PIP}_j = \Pr(\gamma_j = 1 \mid \text{data}),$$

where $\gamma_j = 1$ indicates that gene $j$ is a truly spatially variable gene. In practice, $\text{PIP}_j$ was estimated from the posterior MCMC samples after burn-in as

$$\widehat{\text{PIP}}_j = \frac{1}{M} \sum_{m=1}^{M} \gamma_j^{(m)},$$

where $M$ is the number of retained posterior samples. In this simulation study, we used the last 5000 MCMC iterations after burn-in to estimate the PIP, so $M = 5000$. And $\gamma_j^{(m)}$ is the sampled SVG indicator for gene $j$ at iteration $m$.

A gene was classified as an SVG if its estimated PIP exceeded a predefined threshold $c_{\text{PIP}}$. In our simulation analysis, we set $c_{\text{PIP}} = 0.5$. For each replicate, SVG detection power was then computed as

$$\text{Power} = \frac{\sum_{j=1}^{P} I\left(\gamma_j^{\text{true}} = 1\right) I\left(\widehat{\text{PIP}}_j > c_{\text{PIP}}\right)}{\sum_{j=1}^{P} I\left(\gamma_j^{\text{true}} = 1\right)},$$

where $P$ is the total number of genes and $\gamma_j^{\text{true}}$ denotes the true SVG status used in the simulation.

For each of the six simulation settings, we extracted the replicates containing positively enriched gene sets, that is, $a_{1l} > 0$. Since these positively enriched gene sets are closely aligned with the biological motivation of spaGSE, namely detecting pathways whose member genes exhibit coherent spatial variation in tissue samples. As shown in Figure S3, spaGSE achieved consistently high SVG detection power when $\sigma_\theta = 0.2$, with power tightly concentrated around 98% across different coverage rates. When $\sigma_\theta = 0.5$, the median power decreased to approximately 75-80%, and the variability across replicates increased moderately. This pattern suggests that SVG recovery becomes more challenging when the gene-level spatial summary statistics are noisier, although spaGSE still maintained moderate to high detection power across all coverage rates.

### S5 Sensitivity Analysis

#### S5.1 Sensitivity analysis for the gene-level mixture mean

We further evaluated the sensitivity of spaGSE to the specification of the gene-level mixture mean parameter $\mu_\theta$, which controls the location of the SVG component in the model. In the main simulation study, we used $\mu_\theta = 1$. To assess whether the performance of spaGSE depends strongly on this choice, we considered three alternative values $\mu_\theta \in \{2, 3, 4\}$. The remaining simulation parameters were fixed at CR = 30%, $\sigma_\theta = 0.2$, and $a_{0l} = -2$. As shown in Figure S4, spaGSE maintained stable performance across different choices of $\mu_\theta$. The detection power remained at 99% across all three settings. The FPR was also well controlled, with values close to zero and a maximum observed FPR of 0.11%. These results suggest that spaGSE is robust to the specification of the gene-level mixture mean parameter.

#### S5.2 Sensitivity analysis for the baseline SVG proportion outside of the gene set

We conducted a sensitivity analysis to evaluate the robustness of spaGSE to different choices of the baseline log-odds parameter $a_{0l}$. This parameter controls the background probability that a gene outside the analyzed gene set is spatially variable. We considered four alternative values, $a_{0l} \in \{-0.5, -1, -3, -4\}$, corresponding to expected background SVG proportions of approximately 37.8%, 26.9%, 4.7%, and 1.8%, respectively. In this analysis, the remaining simulation parameters were fixed at CR = 30%, $\sigma_\theta = 0.2$, and $\mu_\theta = 1$. As shown in Figure S5, spaGSE maintained high detection power across all settings, with power remaining between 98% and 99%. The false positive rate was also well controlled, with FPR close to zero across all choices of $a_{0l}$ and a maximum observed FPR of 0.22%. These results suggest that spaGSE is robust to the specification of the baseline SVG proportion.

### S6 Additional Plots for Real SRT Data Analysis

This section presents supplementary heatmaps and spatial patterns for SVGs with clear spatial signals for the real spatial transcriptomics datasets analyzed in Section 5 of the main text. These figures further illustrate pathway-level similarity patterns and support the biological interpretation of spaGSE findings.

#### S6.1 Human Pancreatic Ductal Adenocarcinoma (PDAC) Data

Figure S6 displays the pairwise Pearson correlation heatmap for the 69 significant gene sets identified by spaGSE in the PDAC dataset. The correlation matrix was constructed from the first principal component scores of gene set activity across spatial locations. The heatmap exhibits a clear large-scale block structure. Several identified gene sets are biologically meaningful in the context of PDAC, including ANASTASSIOU_MULTICANCER_INVASIVENESS_SIGNATURE, which reflects tumor invasiveness, SU_HO_CONV_CENT_CHONDROSARCOMA_STROMAL_C4_CANCER_ASSOCIATED_FIBROBLAST, which captures fibroblast-associated stromal programs, JIANG_HYPOXIA_CANCER, which is relevant to hypoxic tumor microenvironments, and KEGG_PANCREATIC_CANCER, which directly represents core pancreatic cancer signaling. Together, these gene sets support the relevance of the detected spatial programs to malignant progression and tumor microenvironment heterogeneity in PDAC.

As shown in Figure S7, the selected 15 SVGs from GRUETZMANN_PANCREATIC_CANCER_UP with set size 81 exhibit highly concordant and spatially localized expression patterns in the PDAC tissue. Since this gene set represents genes commonly up-regulated in pancreatic ductal adenocarcinoma, the observed within-set spatial coherence supports the presence of a shared PDAC-associated molecular signal captured by spaGSE.

As shown in Figure S8, the selected 7 SVGs from NAKAMURA_CANCER_MICROENVIRONMENT_UP with size 13 display coherent spatial expression patterns with a broader and distinct distribution across the tissue. Given that this gene set is related to microenvironment-dependent gene expression, the observed pattern suggests that local tissue context may contribute to the spatial heterogeneity observed in the PDAC sample.

#### S6.2 Human Breast Cancer Data

Figure S9 displays the pairwise Pearson correlation heatmap for the 45 significant gene sets identified by spaGSE in the FFPE breast cancer dataset. The correlation matrix was constructed using the first principal component scores of gene set activity across spatial locations. The heatmap reveals a clear block structure. Several identified gene sets are biologically meaningful in the context of breast cancer, including PUJANA_BREAST_CANCER_WITH_BRCA1_MUTATED_UP, which reflects BRCA1-associated tumor biology, SMID_BREAST_CANCER_LUMINAL_B_DN, which is related to luminal subtype variation, CHARAFE_BREAST_CANCER_BASAL_VS_MESENCHYMAL_UP, which captures basal-to-mesenchymal characteristics linked to tumor classification (Charafe-Jauffret et al. [2006]), and WP_INTEGRATED_BREAST_CANCER_PATHWAY, which represents integrated breast cancer signaling. These signatures contribute to the separation of pathway groups in the heatmap, suggesting that the spatial organization captured by spaGSE reflects coordinated subtype-specific and progression-related programs in the FFPE breast cancer tissue.

As shown in Figure S10, the selected 12 SVGs from CHIN_BREAST_CANCER_COPY_NUMBER_UP with set size 22 exhibit clear within-set spatial coherence in the FFPE breast cancer tissue. Representative genes in this set, including *ERBB2*, *PTK2*, *PAK1*, *NCOA3*, *ZNF217*, and *TNFRSF6B*, show similar localized expression patterns, supporting the interpretation that this pathway captures a shared tumor-associated molecular signal. This spatial concordance is biologically plausible in breast cancer, as amplification-related programs are often closely associated with malignant progression and spatial tumor heterogeneity.

As shown in Figure S11, the selected SVGs from WP_INTERACTIONS_BETWEEN_IMMUNE_CELLS_AND_ MICRORNAS_IN_TUMOR_MICROENVIRONMENT with set size 23 also display coherent spatial expression patterns across the FFPE breast cancer tissue. Representative genes in this set, including *TGFBR2*, *CD80*, *CD86*, *NFKB1*, *NFKB2*, *IRAK4*, *IL4R*, *STAT3*, and *TGFB1* show related localized patterns, suggesting a shared immune- and microenvironment-associated program rather than unrelated individual signals. The observed within-set spatial coherence indicates that spaGSE is able to recover biologically meaningful pathways linked to tumor-microenvironment interactions in breast cancer.

#### S6.3 Human Prostate Cancer Data

Figure S12 displays the pairwise Pearson correlation heatmap for the significant gene sets identified by spaGSE in the prostate cancer dataset. The correlation matrix was constructed from the first principal component scores of gene set activity across spatial locations. The gene sets are clearly clustered into two major groups. One group is enriched for prostate cancer-related and broader cancer signaling signatures, including KEGG_

PROSTATE_CANCER, HWANG_PROSTATE_CANCER_MARKERS, and SETLUR_PROSTATE_CANCER_TMPRSS2_ERG_FUSION_UP. The other group contains gene sets associated with invasion, stromal interaction, fibroblast-related programs, and general cancer progression. This block-wise correlation pattern suggests that the significant gene sets identified by spaGSE are biologically coherent and reflect distinct but coordinated spatial programs in the prostate cancer tissue.

As shown in Figure S13, selected SVGs from KEGG_PROSTATE_CANCER show broadly concordant spatial expression patterns across the FFPE prostate cancer tissue. In particular, the presence of *KLK3*, together with signaling- and proliferation-related genes such as *AKT3* and *CTNNB1*, supports the interpretation that this gene set captures a spatially coherent prostate-cancer-related program. The shared spatial structure of these SVGs is consistent with the biological relevance of KEGG_PROSTATE_CANCER in this dataset.

As shown in Figure S14, selected 8 SVGs from WP_ANDROGEN_RECEPTOR_NETWORK_IN_PROSTATE_CANCER show broadly concordant spatial expression patterns across the FFPE prostate cancer tissue. In particular, androgen-response-related markers such as *KLK3*, *KLK2*, and *FOXA1*, together with regulatory genes including *MYC*, *CDK4*, and *STAT3*, support the interpretation that this gene set captures a spatially coherent androgen-receptor-associated transcriptional program rather than isolated gene-level variation (Alumkal et al. [2020]). The shared spatial structure of these SVGs is consistent with the biological relevance of WP_ANDROGEN_RECEPTOR_NETWORK_IN_PROSTATE_CANCER in this dataset.

#### S6.4 Mouse Embryo Data

Figure S15 displays the pairwise Pearson correlation heatmap for the 50 representative significant gene sets identified by spaGSE in the mouse embryo E16.5 dataset. The correlation matrix was constructed from the first principal component scores of gene set activity across spatial locations. The heatmap exhibits a clear large-scale block structure. Several identified gene sets are directly related to embryonic and organ development, including GOBP_EMBRYO_MORPHOGENESIS, GOBP_EMBRYO_DEVELOPMENT, and GOBP_EMBRYONIC_ORGAN_DEVELOPMENT. In addition, a prominent positively correlated module is enriched for cardiovascular and muscle-related signatures, such as CUI_DEVELOPING_HEART_C2_CARDIOMYOCYTE, KEGG_CARDIAC_MUSCLE_CONTRACTION, and GOBP_MUSCLE_CELL_DIFFERENTIATION. These development- and tissue-specific signatures form broader groups related to embryonic morphogenesis, neurodevelopment, and cardiac/muscle maturation, suggesting that spaGSE captures coordinated spatial variation in developmental programs across different anatomical regions. Together, these results suggest that the gene sets identified by spaGSE are biologically meaningful and reflect clear spatial organization of developmental programs across different embryonic tissues.

As shown in Figure S16, the 9 SVGs in DESCARTES_FETAL_LIVER_HEPATOBLASTS show highly similar spatial expression patterns, with strong and localized enrichment in the annotated liver region. This coherence suggests that the gene set captures a hepatoblast-related developmental program that is spatially organized within the E16.5 mouse embryo. The concordant expression patterns across SVGs further support the biological relevance of this gene set and illustrate that genes within the same functional module can exhibit consistent spatial activity.

As shown in Figure S17, the 22 SVGs in CUI_DEVELOPING_HEART_C2_CARDIOMYOCYTE also display similar spatial expression patterns, with coordinated enrichment in thoracic regions overlapping the annotated heart and nearby muscle-associated structures. This shared spatial pattern is consistent with a cardiomyocyte-related developmental program in the E16.5 mouse embryo. The spatial concordance across SVGs indicates that this gene set represents a coherent developmental module.

### References

Joshi J Alumkal et al. Transcriptional profiling identifies an androgen receptor activity-low, stemness program associated with enzalutamide resistance. *Proceedings of the National Academy of Sciences*, 117(22):12315–12323, 2020.

E Charafe-Jauffret et al. Gene expression profiling of breast cell lines identifies potential new basal markers. *Oncogene*, 25(15):2273–2284, 2006.

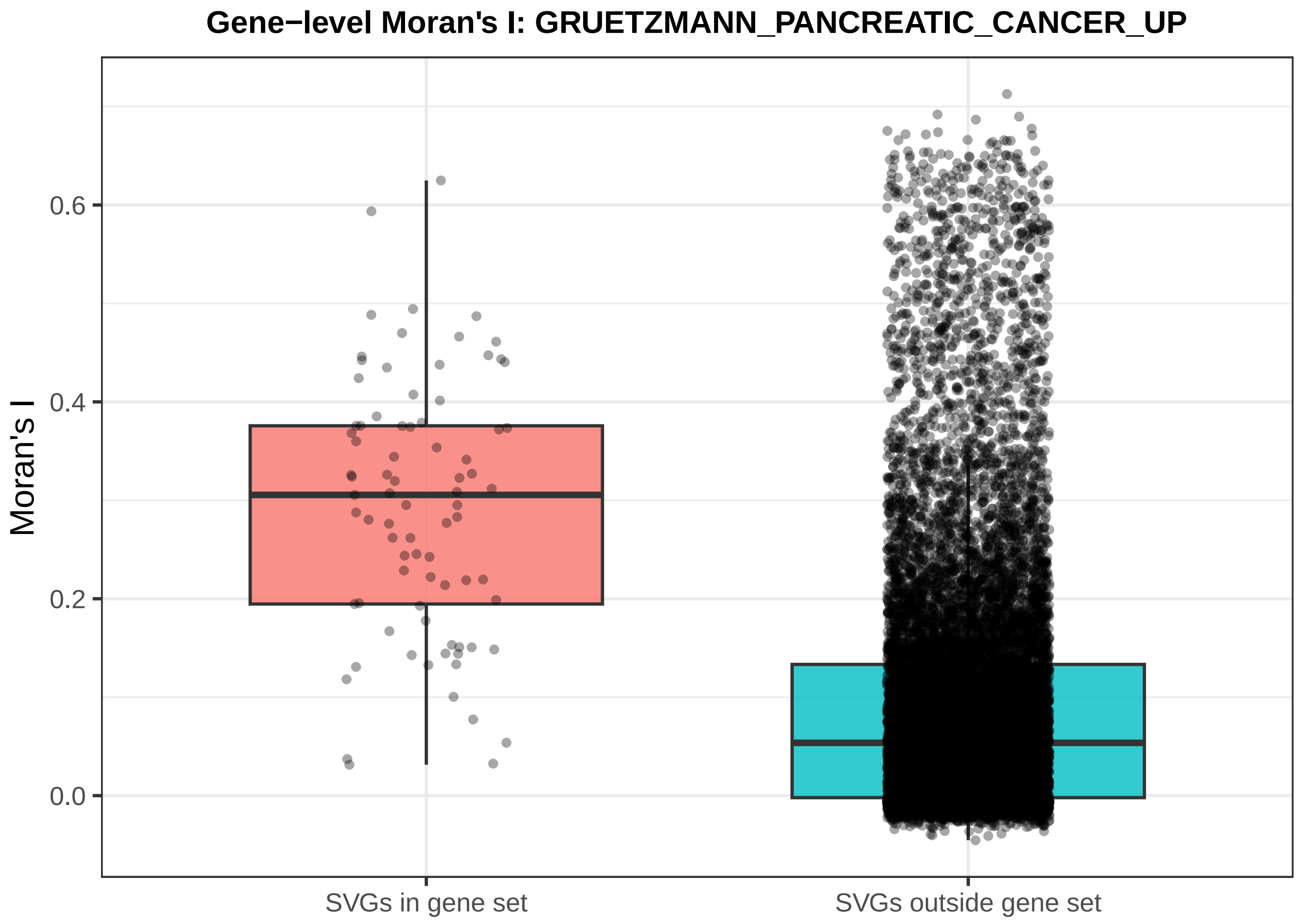


**Figure S1:** Comparison of gene-level spatial autocorrelation between SVGs within and outside the GRUETZMANN_PANCREATIC_CANCER_UP gene set

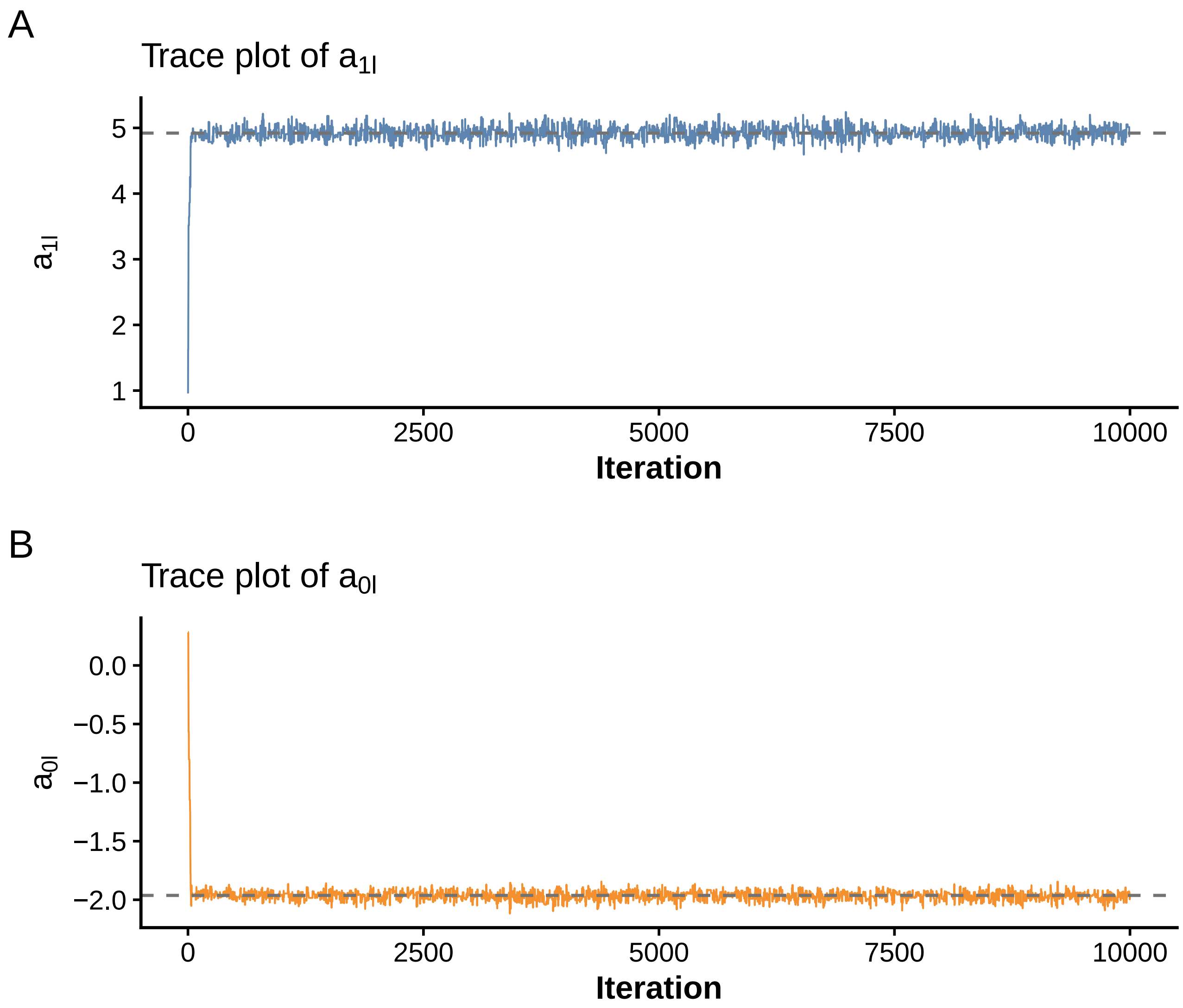


**Figure S2:** Trace plots of gene set enrichment coefficients $a_{0l}$ and $a_{1l}$ under the setting $\sigma_\theta = 0.2, \mathrm{CR} = 30\%$

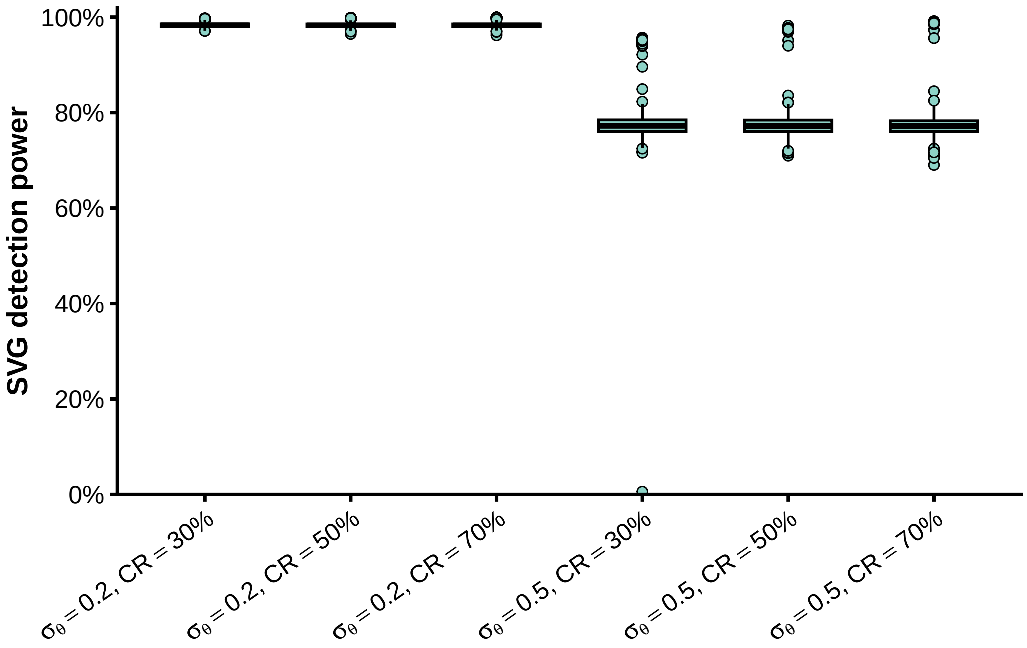


**Figure S3:** Gene-level SVG detection power across six simulation settings. Each boxplot summarizes 100 simulation replicates with either positively or negatively enriched gene sets, that is, $a_{1l} \neq 0$. The six settings vary by the gene-level noise parameter $\sigma_\theta \in \{0.2, 0.5\}$ and the coverage rate CR $\in \{30\%, 50\%, 70\%\}$. For each gene, the posterior inclusion probability was estimated as the posterior mean of the latent SVG indicator $\gamma_j$. SVG detection power was then computed as the proportion of truly spatially variable genes with estimated PIP greater than 0.5.

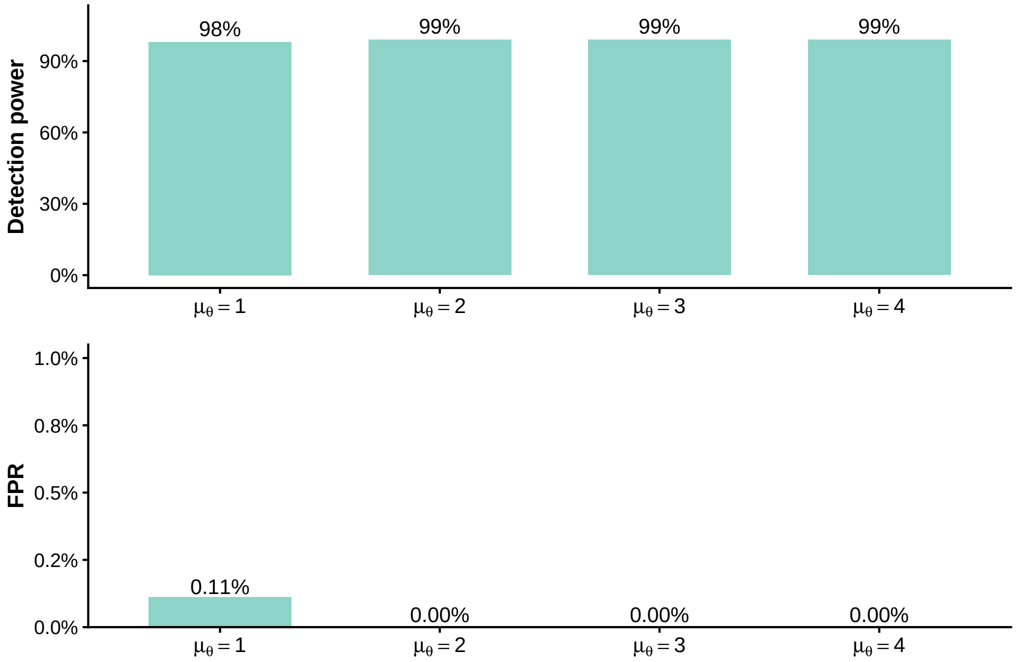


**Figure S4:** Sensitivity analysis of spaGSE with respect to the gene-level mixture mean parameter $\mu_\theta$. The top panel shows detection power, and the bottom panel shows false positive rate (FPR) under different values of $\mu_\theta$. spaGSE achieved consistently high detection power and well-controlled FPR across all considered settings.

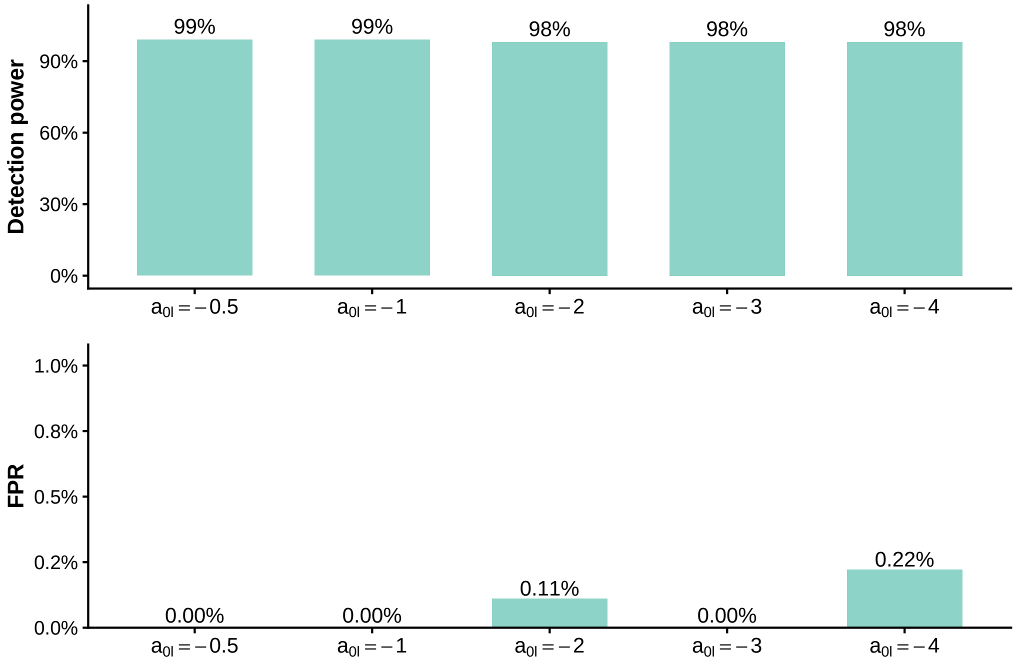


**Figure S5:** Sensitivity analysis of spaGSE with respect to the baseline log-odds parameter $a_{0l}$. The top panel shows detection power, and the bottom panel shows false positive rate (FPR) under different values of $a_{0l}$. spaGSE achieved consistently high detection power and well-controlled FPR across all considered settings.

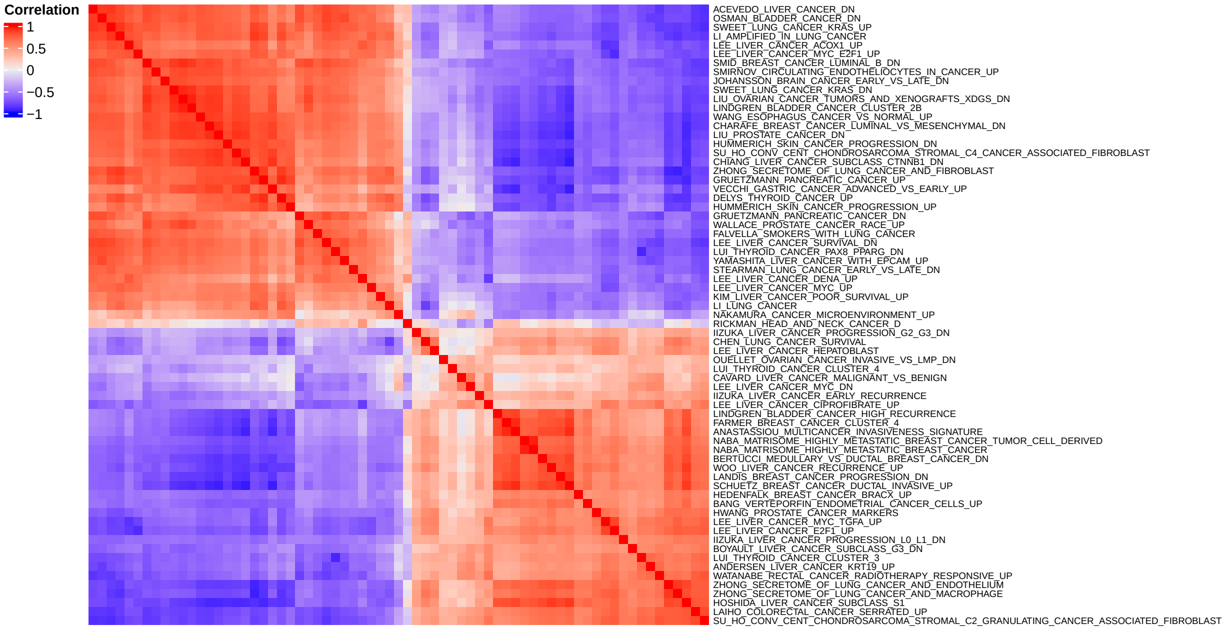


**Figure S6:** A heatmap shows the correlation for 69 significant gene sets identified by spaGSE based on the first principal component after PCA. The gene sets are clearly clustered into two groups

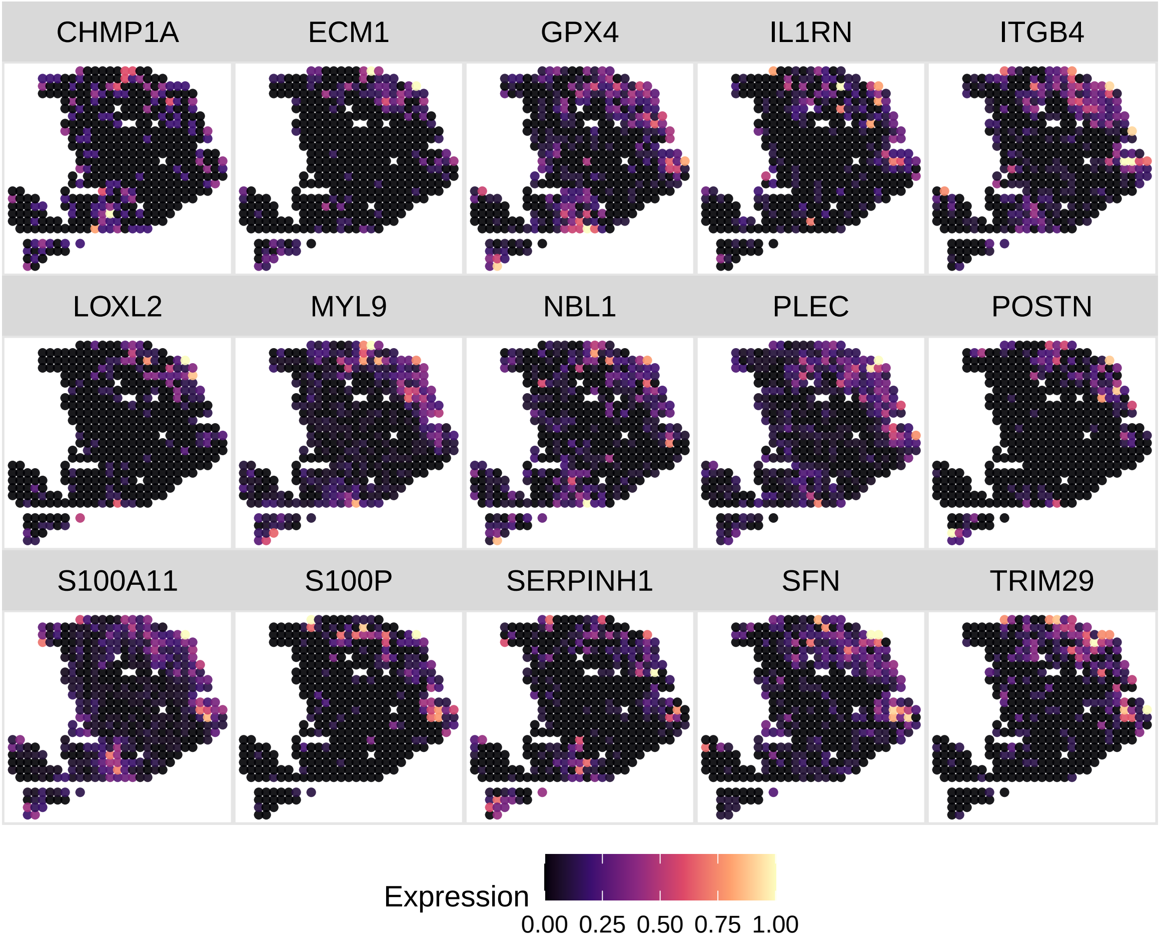


**Figure S7:** Selected 15 SVGs from the gene set `GRUETZMANN_PANCREATIC_CANCER_UP` show highly similar and spatially localized expression patterns, supporting the presence of a coherent PDAC-associated spatial program within this gene set.

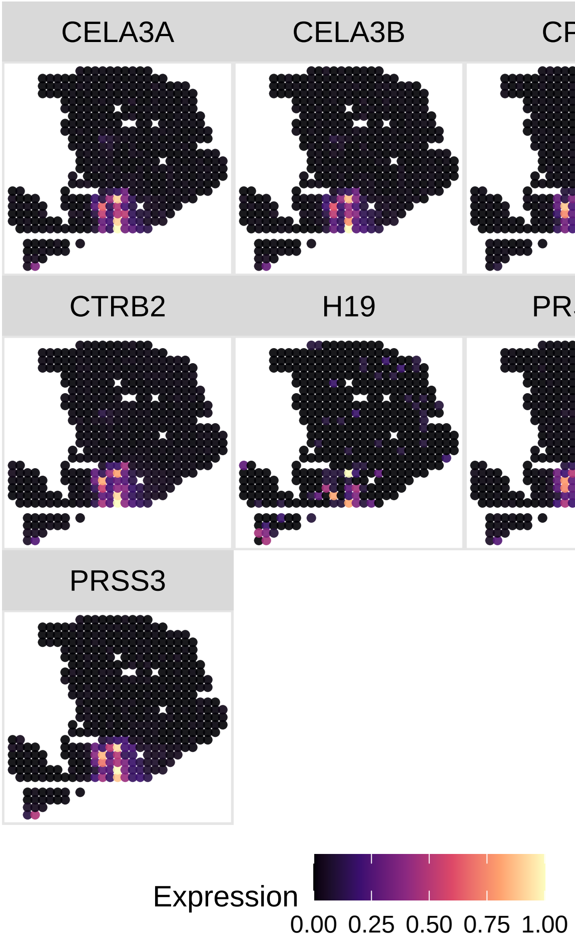


**Figure S8:** Selected 7 SVGs from the gene set `NAKAMURA_CANCER_MICROENVIRONMENT_UP` display coherent spatial expression patterns with a broader distribution across the tissue, consistent with a microenvironment-associated program captured by this pathway.

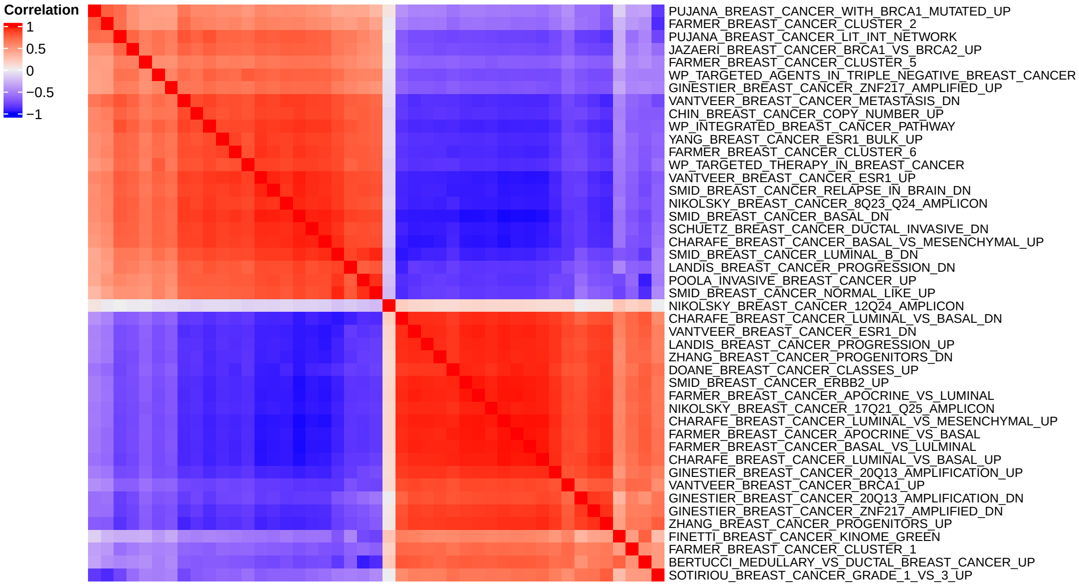


**Figure S9:** A heatmap shows the correlation for 45 significant gene sets identified by spaGSE based on the first principal component after PCA. The gene sets are clearly clustered into two groups

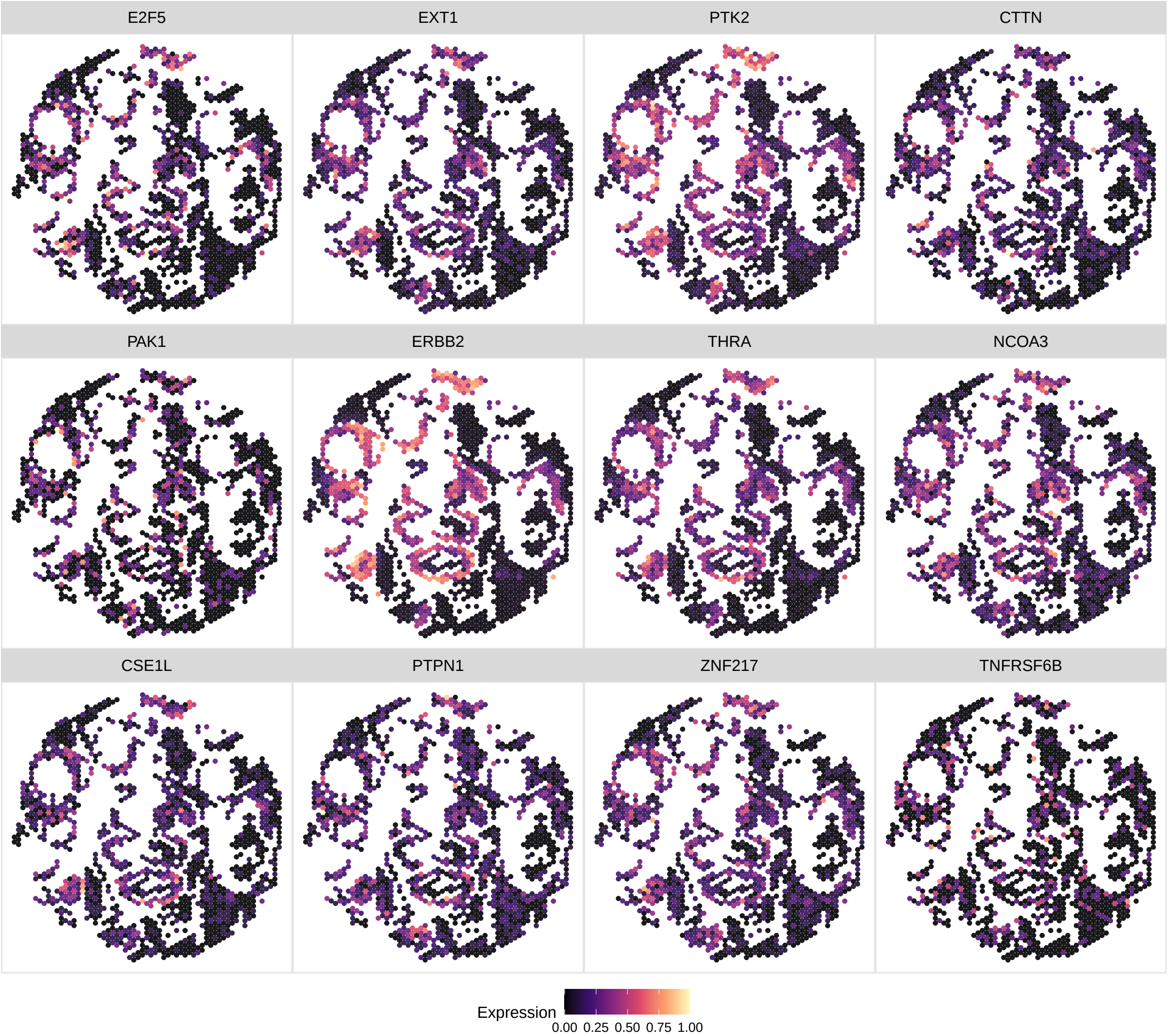


**Figure S10:** Selected 12 SVGs from the gene set `CHIN_BREAST_CANCER_COPY_NUMBER_UP` show coherent spatial expression patterns in the FFPE breast cancer tissue, supporting the presence of a shared tumor-associated molecular signal related to DNA copy number changes.

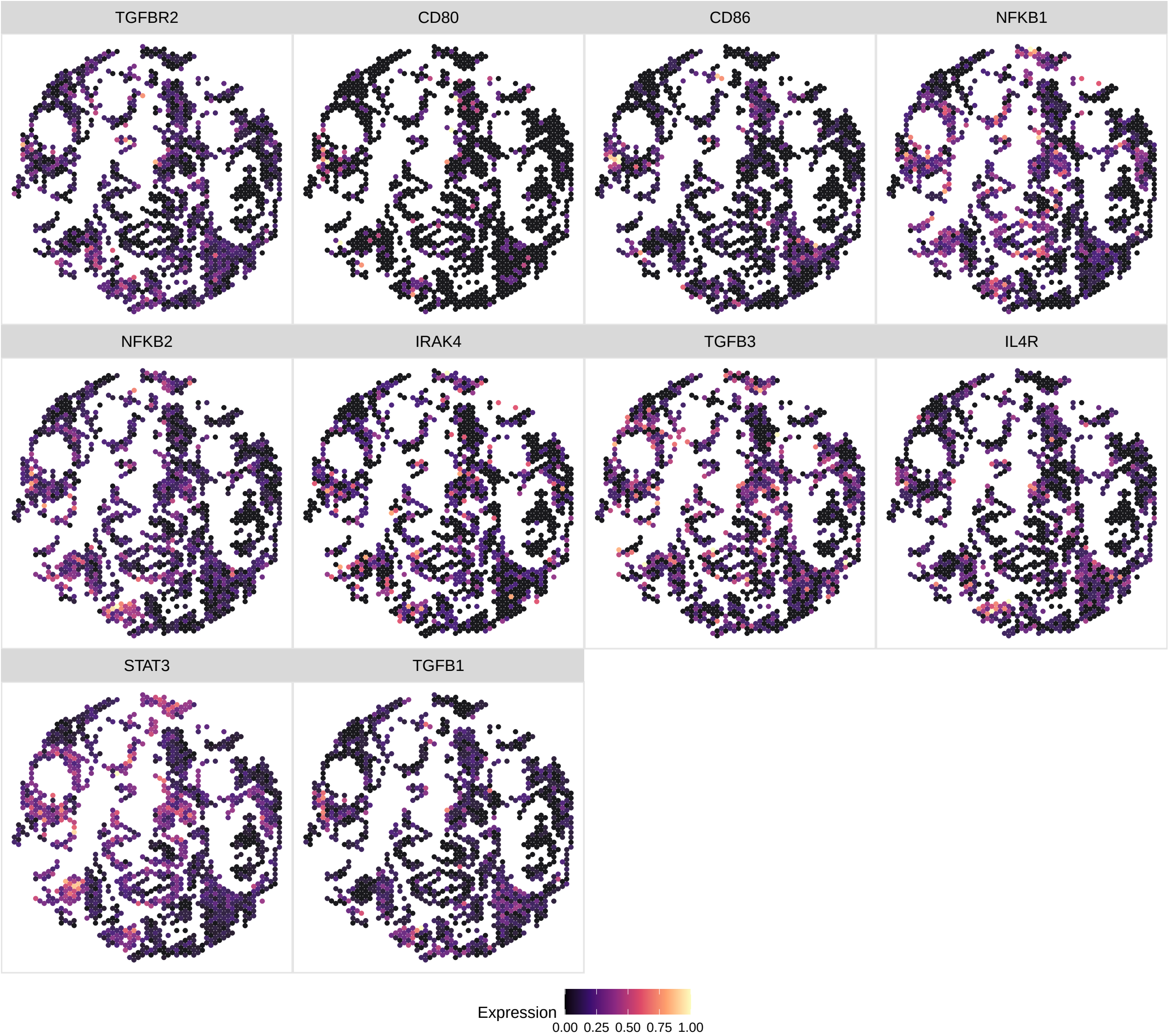


**Figure S11:** Selected 10 SVGs from the gene set `WP_INTERACTIONS_BETWEEN_IMMUNE_CELLS_AND_MICRORNAS_IN_TUMOR_MICROENVIRONMENT` show coherent spatial expression patterns in the FFPE breast cancer tissue, consistent with an immune- and microenvironment-associated program captured by this pathway.

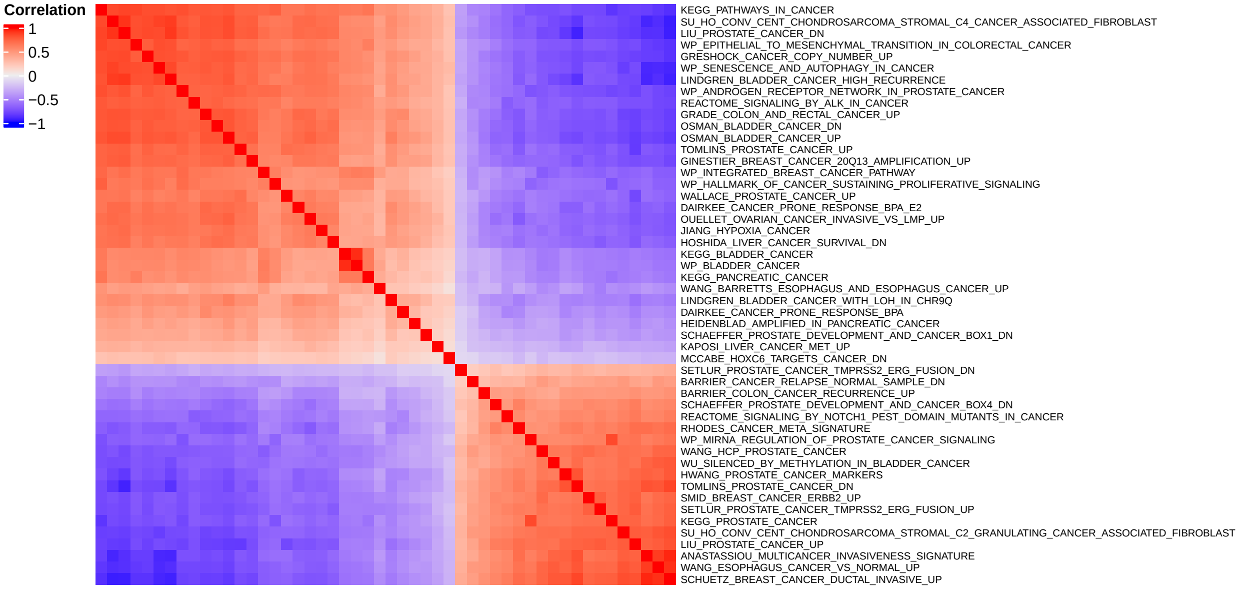


**Figure S12:** A heatmap shows the correlation for representative 50 significant gene sets identified by spaGSE based on the first principal component after PCA. The gene sets are clearly clustered into two groups

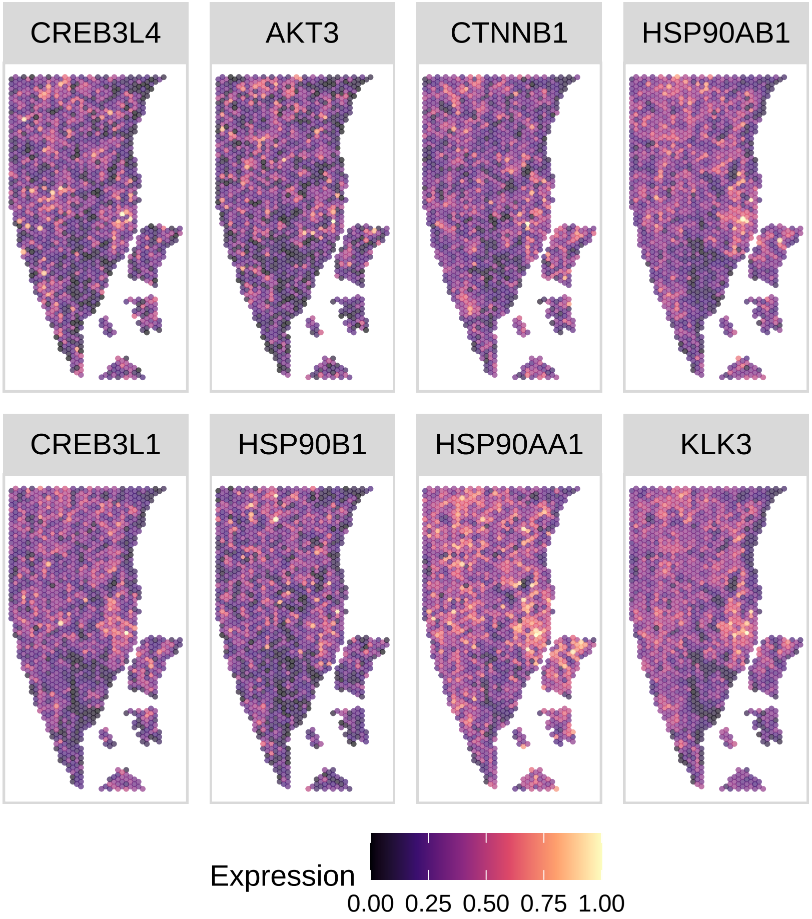


**Figure S13:** Selected 8 SVGs from the gene set `KEGG_PROSTATE_CANCER` show coherent spatial expression patterns in the FFPE prostate cancer tissue, with coordinated enrichment consistent with a prostate-cancer-related transcriptional program.

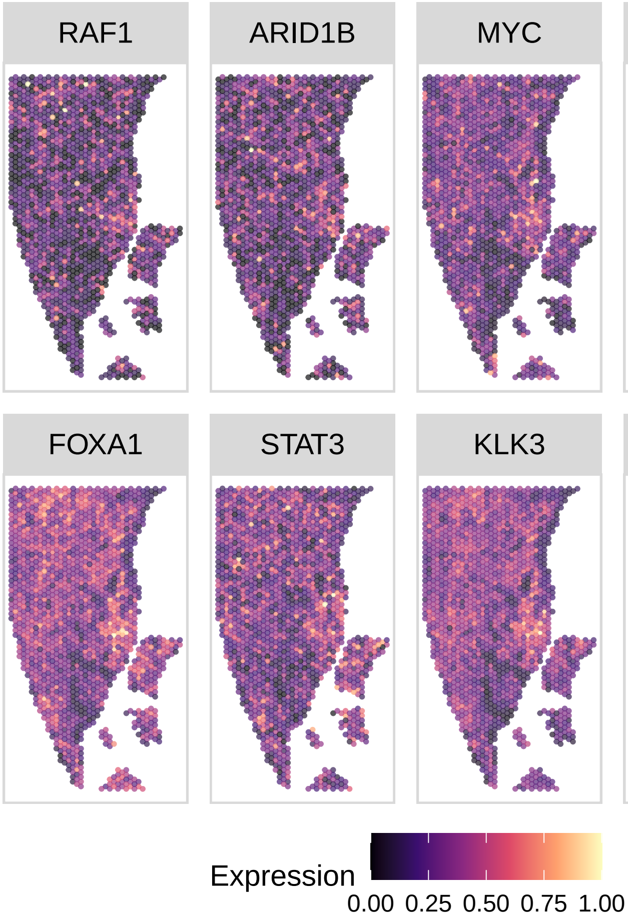


**Figure S14:** Selected 8 SVGs from the gene set `WP_ANDROGEN_RECEPTOR_NETWORK_IN_PROSTATE_CANCER` show coherent spatial expression patterns in the FFPE prostate cancer tissue.

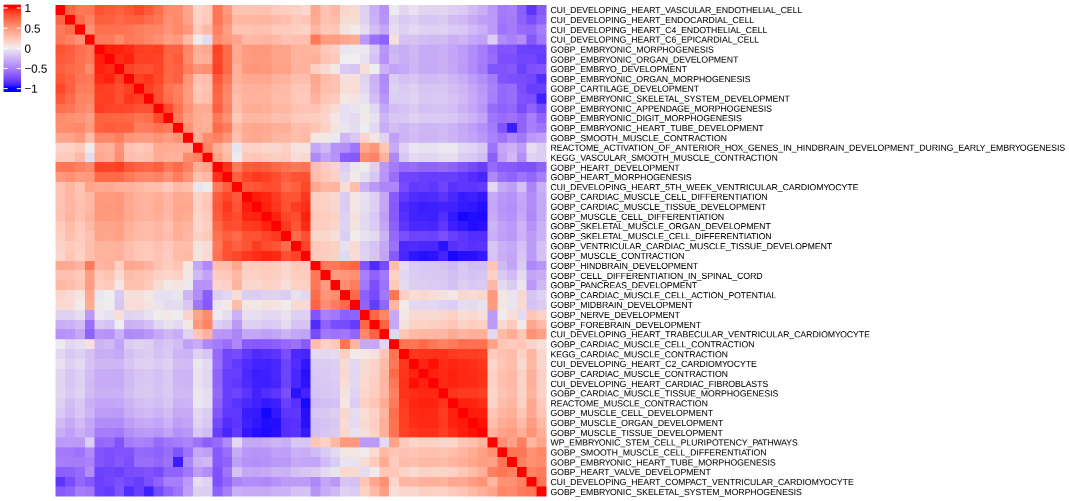


**Figure S15:** A heatmap shows the correlation for 50 representative significant gene sets identified by spaGSE based on the first principal component after PCA. The gene sets are clustered into several groups

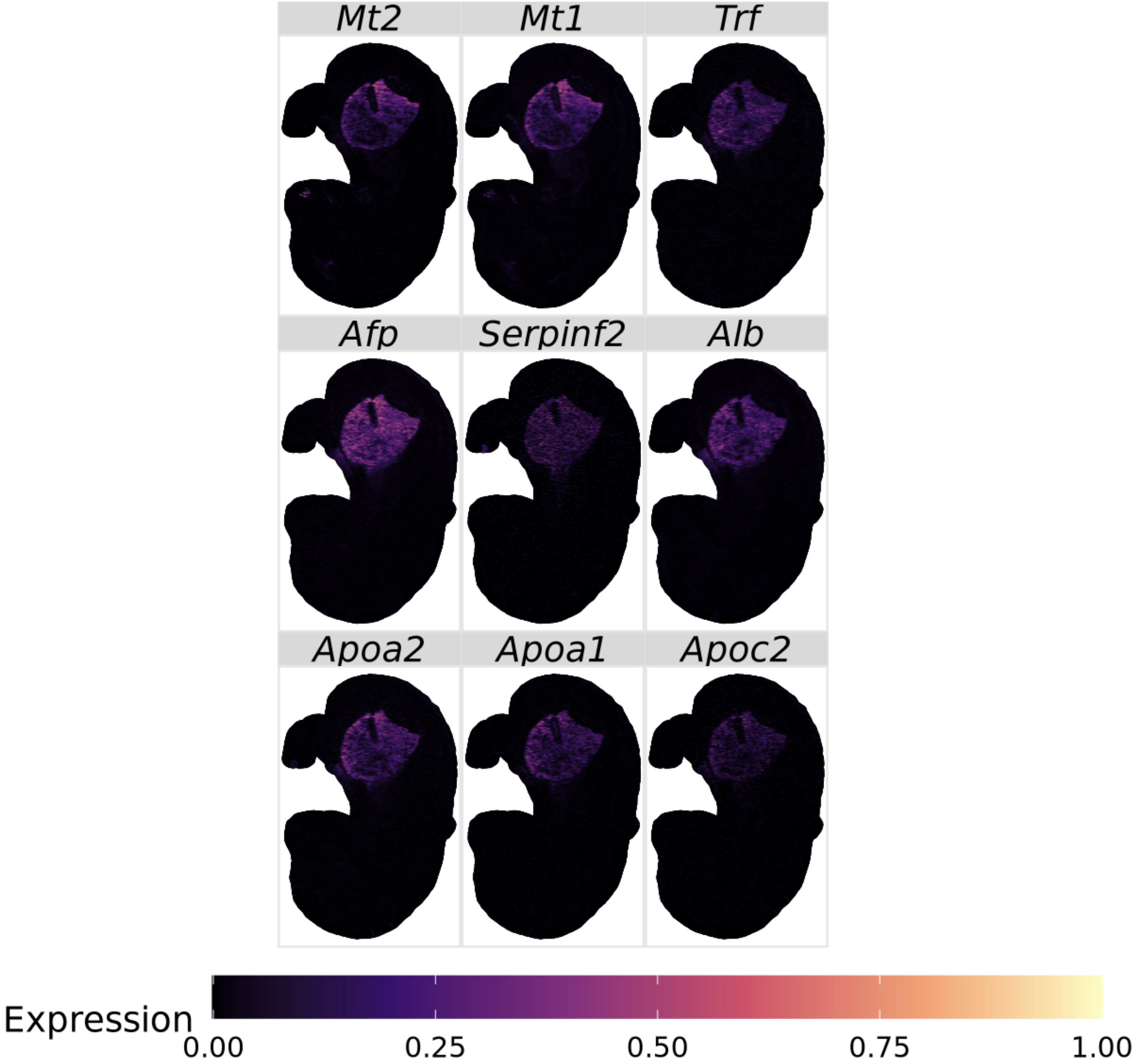


**Figure S16:** Selected 9 SVGs from the gene set DESCARTES_FETAL_LIVER_HEPATOBLASTS show coherent spatial expression patterns in the E16.5 mouse embryo, consistent with a hepatoblast-related developmental program localized to the liver region.

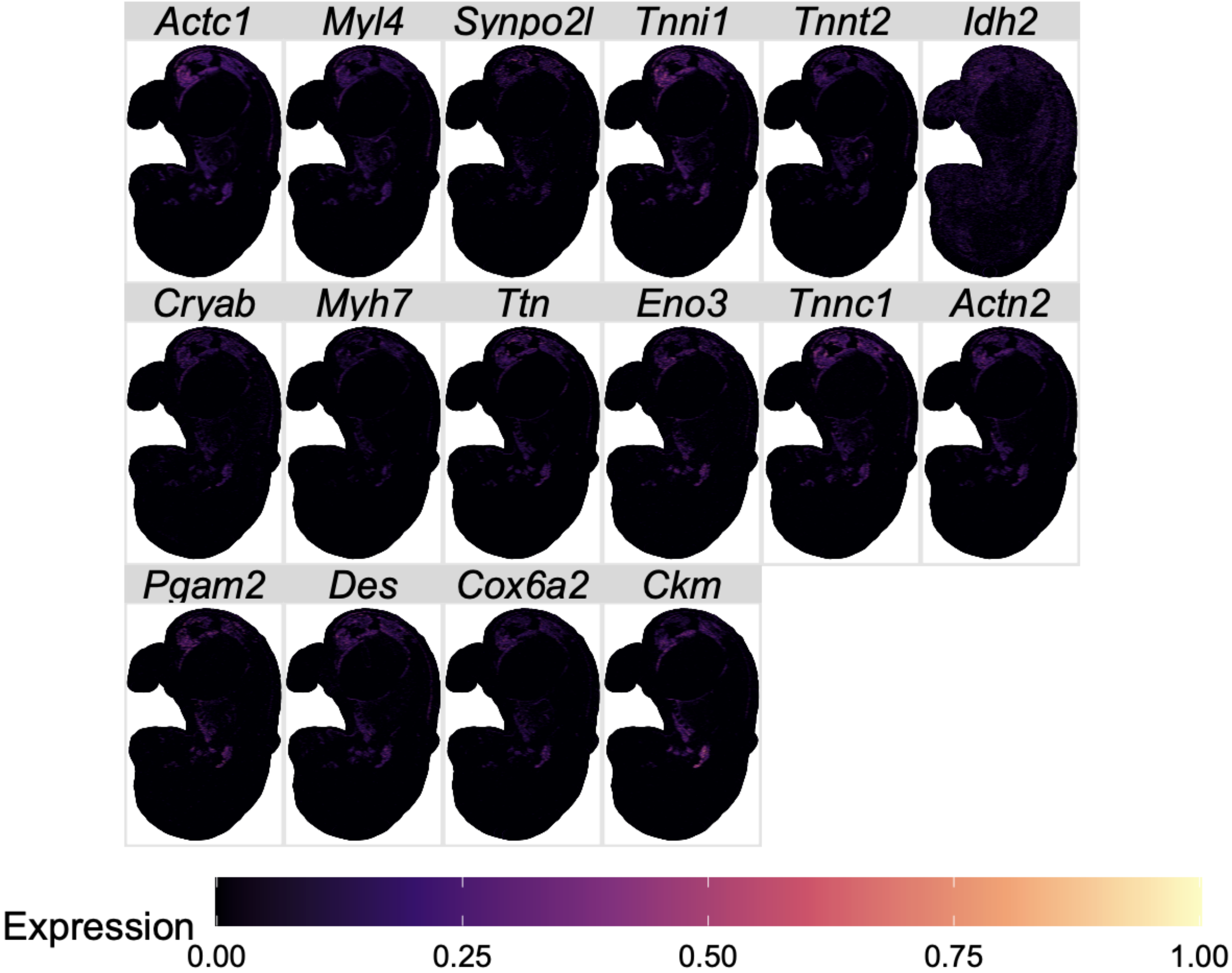


**Figure S17:** Selected 22 SVGs from the gene set `CUI_DEVELOPING_HEART_C2_CARDIOMYOCYTE` show coherent spatial expression patterns in the E16.5 mouse embryo, broadly consistent with a cardiomyocyte-related developmental program.

**Table S1:** The key notations of the proposed spaGSE model.

| | Notation | Support | Definition |
|---|---|---|---|
| Data | $N$ | $N \in \mathbb{N}$ | Number of spatial locations. |
| | $J$ | $J \in \mathbb{N}$ | Number of genes. |
| | $L$ | $L \in \mathbb{N}$ | Number of predefined gene sets. |
| | $\boldsymbol{Y} = [y_{ij}]_{N\times J}$ | $y_{ij} \in \mathbb{N}$ | Gene expression count matrix, where $y_{ij}$ is the count for gene $j$ at spot $i$. |
| | $\boldsymbol{T} = [t_{id}]_{N\times 2}$ | $\boldsymbol{t}_{id} \in \mathbb{R}^2$ | Spatial coordinates of spot $i$. |
| | $\boldsymbol{\theta} = (\theta_1, \ldots, \theta_J)^\top$ | $\theta_j \in \mathbb{R}$ | Gene-level summary statistic measuring spatial variation for gene $j$. |
| | $\boldsymbol{A} = [A_{jl}]_{G\times L}$ | $A_{jl} \in \{0,1\}$ | Gene set annotation matrix, where $A_{jl} = 1$ if gene $j$ belongs to gene set $l$. |
| Latent SVG | $\gamma_j$ | $\gamma_j \in \{0,1\}$ | Latent SVG indicator, where $\gamma_j = 1$ indicates that gene $j$ is spatially variable. |
| | $\pi_j$ | $\pi_j \in (0,1)$ | Probability that gene $j$ is spatially variable. |
| | $\mu_\theta$ | $\mu_\theta \in \mathbb{R}$ | Mean of the spatially variable component in the Gaussian mixture model. |
| | $\sigma^2_\theta$ | $\sigma^2_\theta \in \mathbb{R}^+$ | Variance parameter of the Gaussian mixture components. |
| Enrichment | $a_{0l}$ | $a_{0l} \in \mathbb{R}$ | Baseline log-odds of a gene outside the gene set $l$ being spatially variable. |
| | $a_{1l}$ | $a_{1l} \in \mathbb{R}$ | Enrichment coefficient for gene set $l$; positive values indicate enrichment of SVGs. |
| | $\delta_l$ | $\delta_l \in \{0,1\}$ | Spike-and-slab inclusion indicator for $a_{1l}$. |
| | $\omega_\delta$ | $\omega_\delta \in (0,1)$ | Prior inclusion probability for $\delta_l$. |
| Priors | $\sigma^2_a$ | $\sigma^2_a \in \mathbb{R}^+$ | Slab variance for the enrichment coefficient $a_{1l}$. |
| | $\sigma^2_0$ | $\sigma^2_0 \in \mathbb{R}^+$ | Prior variance for the baseline coefficient $a_{0l}$. |
| | $a_\omega, b_\omega$ | $a_\omega, b_\omega \in \mathbb{R}^+$ | Hyperparameters of the Beta prior for $\omega_\delta$. |
| MCMC | $f(\cdot \mid \cdot)$ | $f(\cdot \mid \cdot) \geq 0$ | Generic likelihood or density function used to evaluate the contribution of observed data under a given latent state or parameter value. |
| | $p(\cdot \mid \cdot)$ | $p(\cdot \mid \cdot) \in [0,1]$ | Generic probability mass or density function for model parameters or latent variables conditional on other quantities. |
| | $J(x;y)$ | $J(x;y) \in [0,1]$ | Proposal probability of proposing state $y$ from the current state $x$ in the Metropolis–Hastings algorithm. |
| | $r_{\text{MH}}$ | $r_{\text{MH}} \in \mathbb{R}^+$ | Metropolis–Hastings acceptance ratio. |
| | $\tau_a$ | $\tau_a \in \mathbb{R}^+$ | Proposal scale parameter for the RWMH update of $a_{1l}$ when $\delta_l = 1$. |
| | $\tau_0$ | $\tau_0 \in \mathbb{R}^+$ | Proposal scale parameter for the RWMH update of $a_{0l}$. |
| Inference | $\text{PIP}_j$ | $\text{PIP}_j \in [0,1]$ | Posterior inclusion probability that gene $j$ is a SVG. |
| | $M$ | $M \in \mathbb{N}$ | Number of retained posterior MCMC samples. |
| | $c_{\text{PIP}}$ | $c_{\text{PIP}} \in [0,1]$ | Threshold used to classify a gene as an SVG. |
| | $I(\cdot)$ | $I(\cdot) \in \{0,1\}$ | Indicator function. |
| | $\text{BF}_{\text{pos}}$ | $\text{BF}_{\text{pos}} \in \mathbb{R}^+$ | Bayes factor measuring evidence for positive enrichment. |
| | $\text{BF}_{\text{neg}}$ | $\text{BF}_{\text{neg}} \in \mathbb{R}^+$ | Bayes factor measuring evidence for negative enrichment. |
